# *Planomonospora*: a Metabolomics Perspective on an Underexplored Actinobacteria Genus

**DOI:** 10.1101/2020.07.19.210815

**Authors:** Mitja M. Zdouc, Marianna Iorio, Sonia I. Maffioli, Max Crüsemann, Stefano Donadio, Margherita Sosio

## Abstract

Despite an excellent track record, microbial drug discovery suffers from high rates of re-discovery. Better workflows for the rapid investigation of complex extracts are needed to increase throughput and allow early prioritization of samples. In addition, systematic characterization of poorly explored strains is seldomly performed. Here, we report a metabolomic study of 72 isolates belonging to the rare actinomycete genus *Planomonospora*, using a workflow of open access tools to investigate its secondary metabolites. The results reveal a correlation of chemical diversity and strain phylogeny, with classes of metabolites exclusive to certain phylogroups. We were able to identify previously reported *Planomonospora* metabolites, including the ureylene-containing oligopeptide antipain, the thiopeptide siomycin including new congeners and the ribosomally synthesized peptides sphaericin and lantibiotic 97518. In addition, we found that *Planomonospora* strains can produce the siderophore desferrioxamine or a salinichelin-like peptide. Analysis of the genomes of three newly sequenced strains led to the detection of 47 gene cluster families, of which several were connected to products found by LC-MS/MS profiling. This study demonstrates the value of metabolomic studies to investigate poorly explored taxa and provides a first picture of the biosynthetic capabilities of the genus *Planomonospora*.

## Introduction

Natural products are excellent sources for bioactive scaffolds. Over the last four decades, 66% of approved small-molecule drugs were actual natural products or at least inspired from such.^1^ This is an impressive track record, considering the general withdrawal of industrial activity from the field.^2^ One reason for this disinterest has been the frequent rediscovery of known molecules in activity-based screenings, especially in microbe-derived extracts.^3^ However, the focus on bioactivity as selection criterion provides a biased perspective on a small portion of the chemical diversity microbes are capable to produce. Despite decades of research, the majority of secondary metabolites remain “metabolomic dark matter”,^4^ with high probability of structural novelty^5^ and novel bioactive scaffolds.^6^ Still, streamlined approaches for strain prioritization and workflow optimization are needed to render drug discovery from microbial sources a cost-effective endeavor. ^5^

Recent advances in genome mining have enabled researchers to investigate the biosynthetic potential of bacteria *in silico*, with minimal wet lab work.^7^ Tools such as antiSMASH^8^ allow to mine genomes for biosynthetic gene clusters (BGC), while BGC repositories such as MIBiG^9^ aid in the evaluation of BGC novelty. In addition, advances in (tandem) mass spectrometry and the introduction of molecular networking,^10^ the tandem mass (MS^2^) based grouping of molecules by structural relatedness, has made untargeted metabolomics broadly available,^11^ while public databases in the likes of GNPS^12^ and the Natural Product Atlas^13^ facilitate metabolite annotation. These methods allow to rationalize resources and quickly prioritize strains or metabolites for further investigations. Earlier studies on bacterial genera, such as the actinobacteria *Salinispora*^14–16^ and *Nocardia*,^17^ the myxobacterium *Myxococcus*^18^ and the gamma-proteobacterium *Pseudoalteromonas*^19^ have demonstrated distinct chemical profiles and shown correlations between taxonomic and metabolomic diversity.

We are particularly interested in exploring the metabolic capabilities of “rare” genera of actinomycetes present within the Naicons collection, which comprises approximately 45,000 actinomycete strains of diverse origin, isolated between 1960 and 2005.^20^ One such genus is *Planomonospora*. Originally described by researchers from Lepetit (the predecessor company of Naicons) in 1967, ^21^ so far, just six species, two sub-species and four unclassified strains can be found in public collections or databases. Only few molecules have been described as produced by representatives of this genus: the thiopeptides thiostrepton ^22^ and sporangiomycin, ^23^ the latter identical to siomycin, produced also by *Streptomyces sioyaensis*^24^ (and henceforth referred to as such); lantibiotic 97518, also known as planosporicin,^25,26^ a member of a family of class I lantipeptides produced by many actinobacterial genera; ^27^ the lassopeptide sphaericin; ^28^ and the ureylene-containing oligopeptide antipain, ^29^ which is also produced by *Streptomyces*. ^30^ Here, we aim to assess the capability of *Planomonospora* strains to produce secondary metabolites, using a pipeline of freely available tools for metabolome and genome mining. The modular nature of our workflow facilitates fast and flexible processing of data, allowing to quickly prioritize strains and/or metabolites for further investigation (see Figure 1). The investigation was carried out on 72 strains from the Naicons collection and was complemented by selected genomic analyses. This study gives unprecedented insight into the rare genus *Planomonospora*, correlating metabolites to their putative biosynthetic gene clusters and making way for targeted isolation efforts.

**Figure 1.**
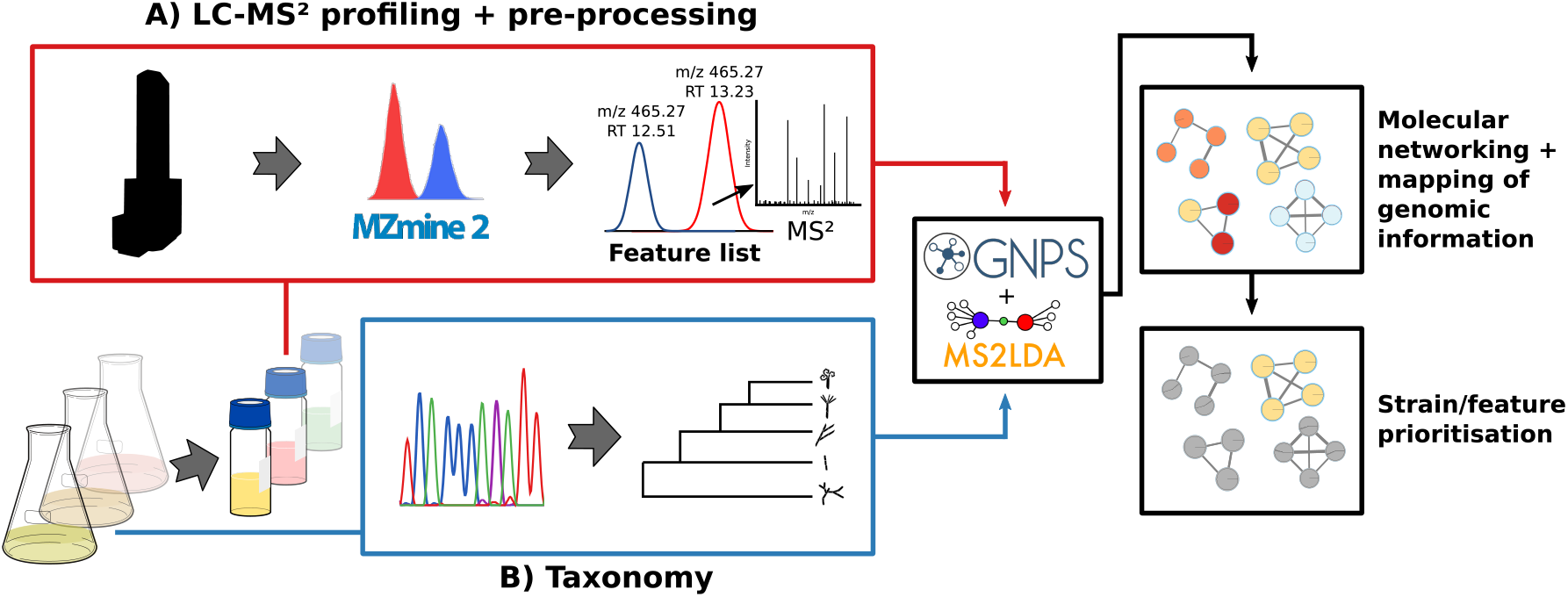
Workflow: Strains are cultivated, extracts are prepared and analyzed. (A) Data dependent acquisition (DDA) mode on an ESI-HR-LCMS-MS-instrument. Data are preprocessed with the MZmine2 software, yielding a list of features, which are then analyzed by GNPS feature-based molecular networking, with consecutive MS2LDA curation. (B) Taxonomy of the strains is established by 16S rRNA-sequencing. Strains/features are prioritized for consecutive targeted isolation.

## Results and Discussion

### Determination of Phylogenetic Affiliation

From the over 350 *Planomonospora* entries listed in the Naicons collection, we selected 72 strains confirmed by 16S rRNA gene sequencing as belonging to the genus *Planomonospora*. As far as information was available, all strains were isolated from soil samples, with the majority originating from central Africa and the Mediterranean region (see Figures S1 and S2). Overall, the 72 strains yielded 35 unique 16S rRNA sequences, 31 of which were not reported previously (see Figures S1 and S2). The resulting phylogenetic tree (see Figure 2) was found to be in agreement with previous studies^31,32^ and showed three phylogroups with a relevant number of representatives: phylogroup “C” includes 13 strains (9 of them from Naicons collection) and 9 distinct 16S rRNA sequences; phylogroup “A2” includes 12 strains (11 of them from Naicons collection) and 8 distinct 16S rRNA sequences; and the most populated phylogroup “S” includes 46 strains (43 of them from Naicons collection) and 13 distinct 16S rRNA sequences. In addition, the phylogenetic analysis yielded three poorly represented phylogroups: “V1”, which includes *Planomonospora venezuelensis* JCM3167 and Naicons strain ID43178, with identical 16S rRNA sequences; the somehow related “V2” group, with six distinct Naicons isolates with identical sequences; and “A1”, with just two Naicons strains with identical sequences. All phylogroups contained sequences of previously described *Planomonospora* species, except “V2” and “A1”. Given the extent of sequence distance from validly described species (see Figures S1 and S2), many of the Naicons strains likely represent new species within this genus. In the following analyses, we consider “V1” as a phylogroup, even though it contains only one strain.

**Figure 2.**
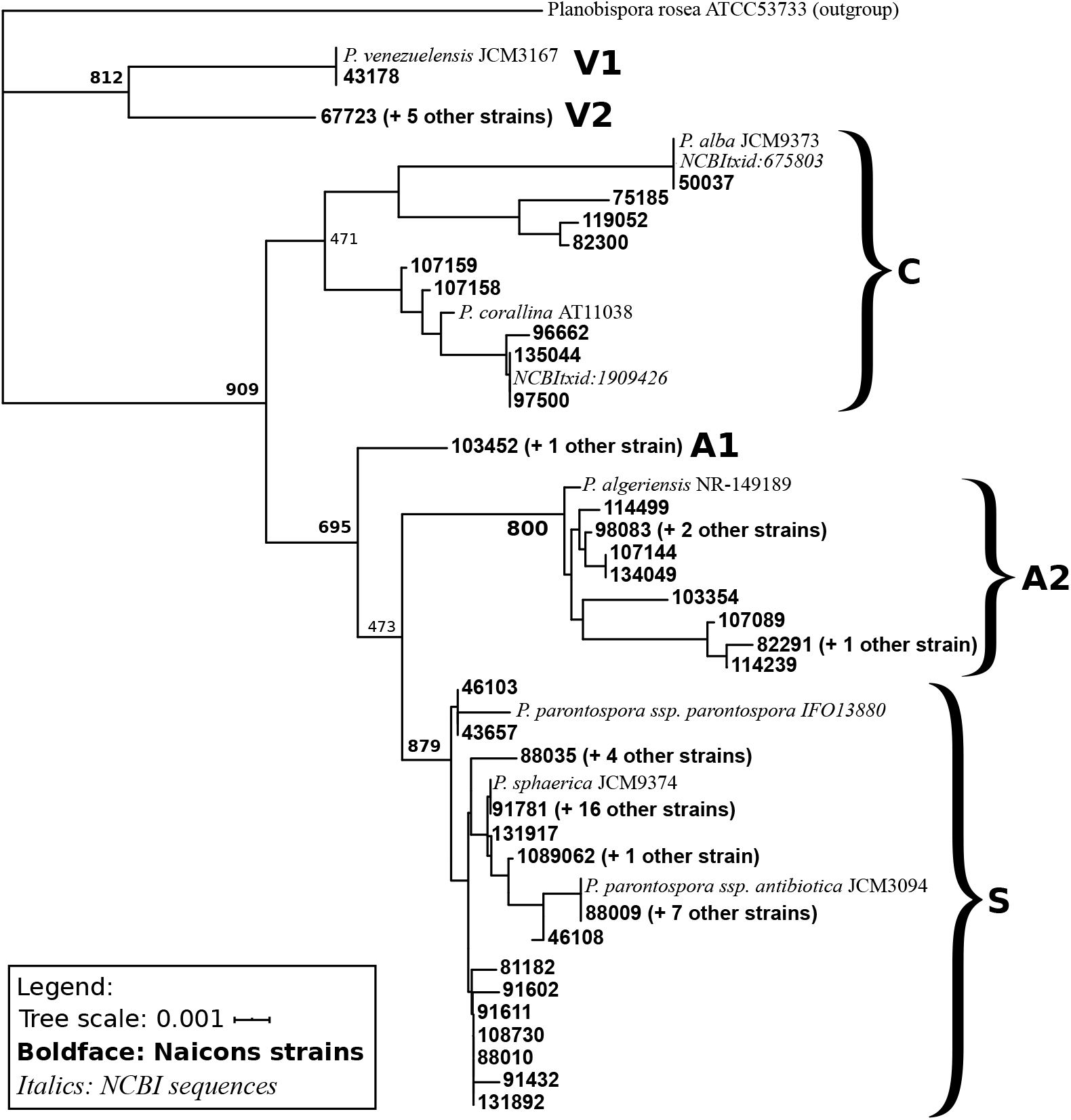
16S rRNA-based phylogenetic tree of *Planomonospora(P.)-strains*. Naicons strains are represented by their ID numbers in boldface. Type strains and two unclassified strains with complete 16S sequences are indicated in italics. Naicons strains with identical sequences are represented by a single ID number, with the number of additional strains in parentheses (details in Figures S1 and S2). Bootstrap values (1000 times resampled) higher than 60% are indicated in bold type. *Planobispora rosea* ATCC53733 was used as outgroup.

### Cultivation, Extraction and Molecular Network Analysis

To find appropriate cultivation conditions for the *Planomonospora* strains, we first explored the behavior of a selected number of isolates under a variety of different conditions, including three solid and six liquid media. Four liquid media (AF, R3, MC1 and AF2, see Experimental Procedures) afforded the highest metabolic diversity (data not shown) and were used in the following analyses.

Cultivation of the 72 strains in the four media and preparation of two extracts per culture yielded 576 samples. To expedite analysis, the two extract from each culture were combined, yielding 288 samples, two of which were removed due to cross-contamination. The remaining 286 samples were analyzed by high resolution ESI-LC-MS/MS in data dependent acquisition mode. The LC-MS/MS data were subjected to a workflow consisting of several steps (see Figure 1): files were (i) preprocessed with the feature finding tool MZmine2 to take into account chromatography well-resolved isomers as well as to correct for *m/z* and retention time drift; ^33,34^ (ii) analyzed using the feature-based molecular networking workflow of GNPS;^12,35^ and (iii) visualized using the program Cytoscape^36^ (see Figure S3).

As seen in Figure 3, the molecular network contains 1492 features, with media components, background impurities (from the extraction process) as well as features with less than *m/z* 300.0 removed beforehand (to exclude primary metabolism). Of those 1492 features, 447 (30%) were organized in 60 molecular families. The remaining 1045 features were singletons. The molecular network is a visual representation of the chemistry detected by mass spectrometry. The network is comprised of sub-networks (called molecular families), which are in turn made up of features (=nodes) and edges. Each feature is a non-redundant representation of a detected parent ion over all samples, with a distinct mass-to-charge-ratio *(m/z)*, retention time (RT) and corresponding tandem mass (MS^2^) fragmentation spectra (see Figure S3). For example, soyasaponin I, a soymeal-related medium component, would be represented as a single feature with a specific *m/z*, RT and MS^2^ spectrum, even though it is detected in almost all samples. Still, the number of features is not equivalent to the number of metabolites, since the same metabolite can be detected as different adducts (and thus features) by ESI mass spectrometry (e.g. [M+H]^+^, [M+2H]^2+^, [M-2H+Fe] ^+^, [M+Na]^+^ …). Features can be further connected by edges, which represent the pairwise similarity between two MS^2^ fragmentation spectra. Since similar MS^2^ fragmentations indicate a similar chemical structure of the parent ions, connected features are considered to be structurally related. ^10^ Molecular families are therefore topological representations of chemical relatedness. Features can also be singletons, if they have sufficiently unique MS^2^ spectra not to cluster with any other feature. For the following investigation, all 1492 features were taken into account, independent of their topological organization (singleton or member of a molecular family).

**Figure 3.**
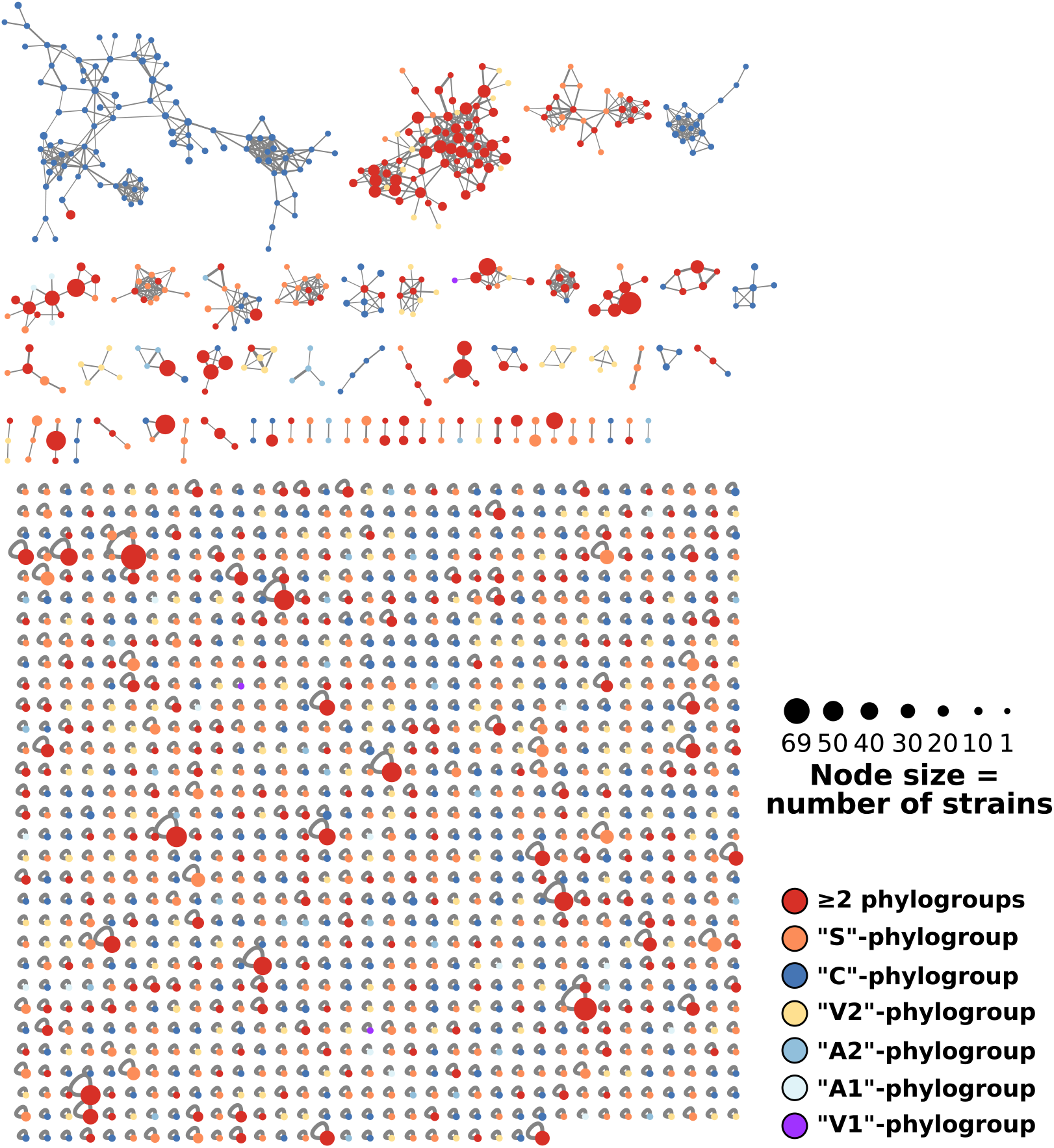
Complete molecular network of 286 *Planomonospora* extracts, encompassing 1492 features (=nodes). 447 features were organized in 60 molecular families. Node size correlates to the number of contributing strains, while the colors give the contributing phylogroup(s).

To explore trends in feature distribution, sample metadata, such as producing strain, cultivation medium and phylogroup affiliation, can be mapped onto the molecular network, which supports intuitive assessment and helps in data organization. An example is shown in Figure 3, where node color correlates to phylogroups. After this sort of visualization, we systematically explored occurrence of features according to strain, phylogroup and medium, as explained below.

We first investigated how features varied within and between phylogroups. A recent study on myxobacteria demonstrated strong correlation between taxonomic and secondary metabolite diversity, i.e. metabolite profiles showed high taxonomic specificity. ^37^ This raised the question whether this applied to *Planomonospora* as well. Indeed, only 1% of features were detected in members of all phylogroups, as shown by the Venn diagram in Figure 4A. The vast majority (74%) of the 1492 features were phylogroup-specific, meaning that they were not detected in samples derived from strains of a different phylogroup. The number of specific features was especially high for phylogroup “C”. Despite counting only 9 strains, 31% of all detected features were exclusive to its members. This is consistent with the phylogenetic tree of Figure 2, which indicates that phylogroup “C” is more divergent from the other well-represented phylogroups “A” and “S”. In contrast, the number of phylogroup-specific features was relatively low in groups “A1” and “A2”, suggesting that the separation into two phylogroups might be an artifact due to the existence of only one 16S rRNA sequence in phylogroup A1. Overall, the results suggest that secondary metabolite production in *Planomonospora* ssp. is a phylogroup-defining trait.

**Figure 4.**
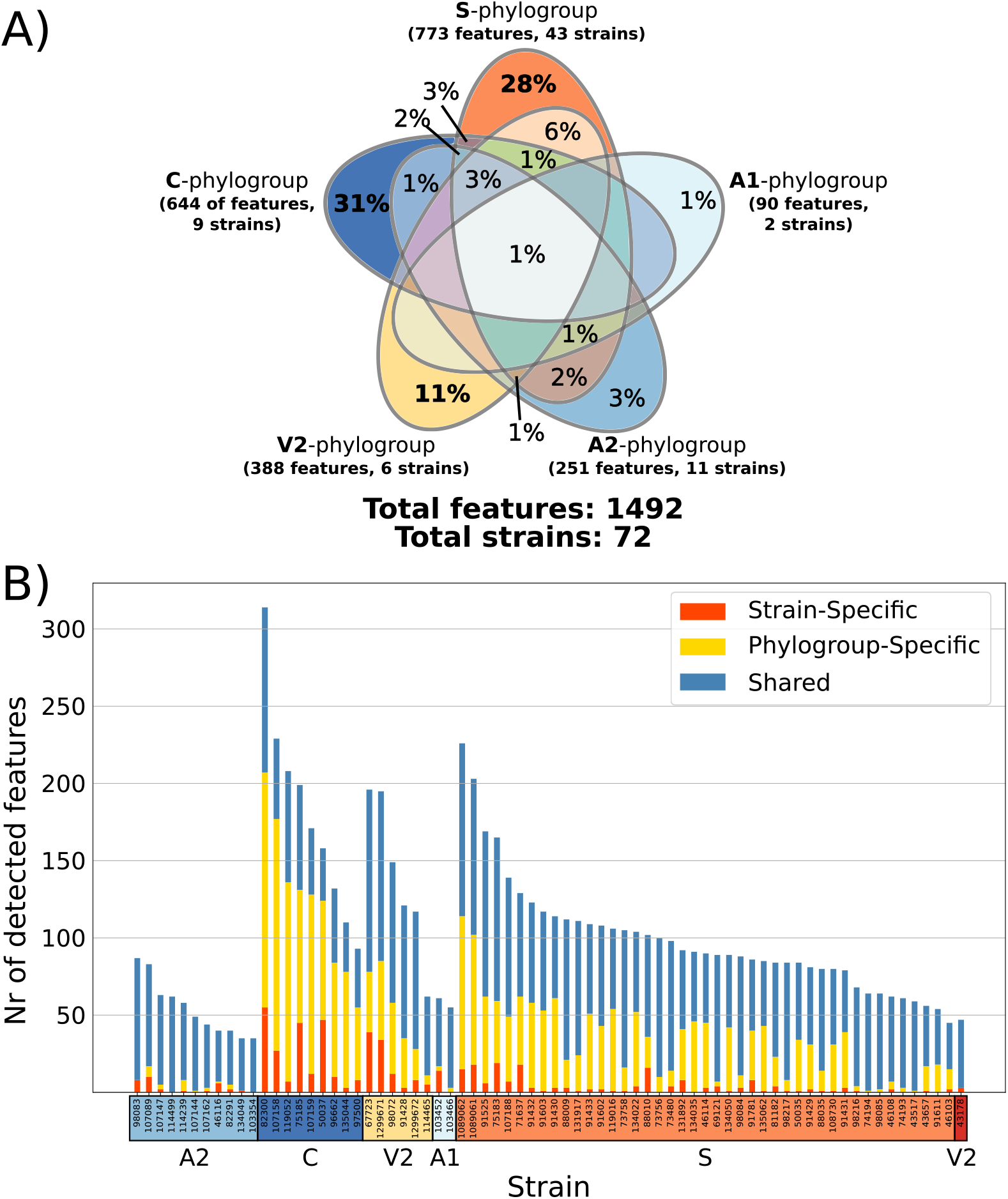
Distribution of 1492 features according to (A) phylogroups and (B) strains. In panel A, overlaps amounting to less than 1% are not labeled, while features detected in phylogroup “V1” were omitted from the analysis. In panel B, each bar represents a different strain. Bars are separated into strain-specific (red), phylogroup-specific (yellow, detected in at least one additional strain from the same phylogroup) and shared features (blue, detected in at least one additional strain from a different phylogroup).

We also investigated the distribution of features at a strain level. Each bar in Figure 4B represents a single strain, with the number of detected features indicated on the y-axis. It can be seen that this number varies greatly among strains, with some “talented” strains standing out in terms of both total features and strain-specific features. Phylogroup “C” again, and, to a lesser extent, “V2”, were enriched in such strains. This is particularly relevant for phylogroup “V2”, for which all six strains shared an identical 16S rRNA sequence. In total, 36% of features were strain-specific, indicating that also for the genus *Planomonospora*, secondary metabolites tend to be strain-specific.

We were also interested in the effects of the different cultivation media on metabolite profiles of strains. Since all 72 strains were cultivated in the same four media, feature distributions could be evaluated (see Figure 5). Overall, 57% of features were medium-specific. Media MC1 and R3, with 26 and 18% of exclusive features, respectively, were the biggest contributors. Further, these two media covered 85% of all detected features. In contrast, just 9% of features were found in all media. Similar observation were made in a study of 26 marine *Streptomyces* strains, in which 71% of detected ions were medium-specific and just 7% common to all 3 conditions.^14^ Even though complete metabolite coverage remains elusive, our results suggest that two media should cover most of the metabolites produced by different *Planomonospora* strains belonging to different phylogroups.

**Figure 5.**
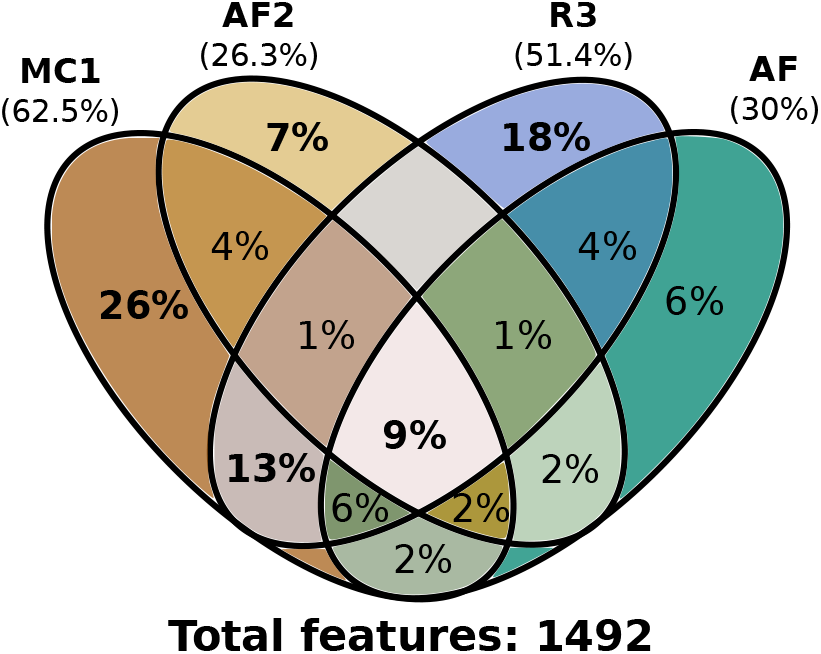
Visualization of the distribution of features in extracts with regard to cultivation medium. Media are MC1, AF2, R3 and AF. Overlaps amounting to less than 1% are not labeled.

### Metabolite Annotation

Next, we started examining the actual metabolites present in the *Planomonospora* metabolome by first identifying known metabolites in a process termed de-replication.^38,39^ In addition to providing insights into the biosynthetic potential of this poorly studied genus, establishing the known metabolites can highlight features likely to be associated with novel chemistry, thus evading the pitfalls of re-investigating reported compounds. As described below, we de-replicated representatives of 11 molecular families, some of which are visualized in Figure 6. For annotation, we used both spectral matching (comparison with identified spectra in curated databases, such as GNPS) and literature search. To increase confidence in the annotations, Chemical Analysis Working Group (CAWG) criteria^40^ were applied, leading to a number of class 1 (comparison against an authentic standard) and class 2 (putatively identified molecule) annotations (for a full list, see Figure S4).

**Figure 6.**
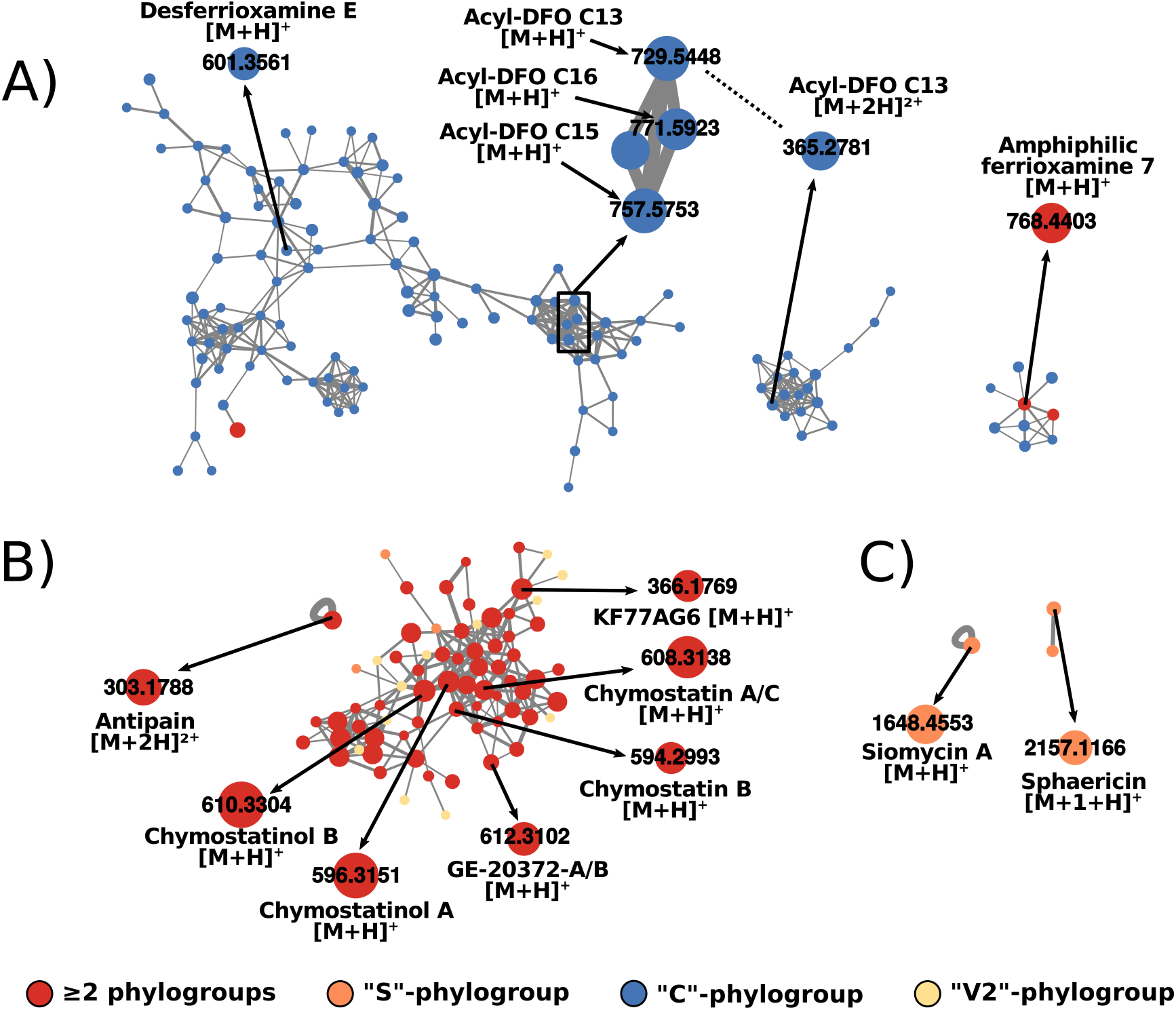
Visualisation of selected annotated molecular families, with features annotated manually or by GNPS spectral library search (e.g. “amphiphilic ferrioxamine 7”). (A) desferrioxamines (DFO) are mostly occurring in strains from phylogroup “C”. (B) chymostatin-like metabolites show no phylogroup-specificity. (C) Siomycin and sphaericin are exclusive to phylogroup “S”.

We first searched for the metabolites previously reported from *Planomonospora*. The lantibiotic 97518 was identified in an extract from strain ID50037 after observing a signal with *m/z* 1097.39, which matched the [M+2H]^2+^ ion (representing the isotope containing 1 ^13^C atom), and comparison of its MS^2^ fragmentation spectrum to the one reported in the literature confirmed this assumption (see Figures S4 and S31).^25^ The lassopeptide sphaericin, identified by the signal *m/z* 2157.12, corresponding to the [M+H]^+^ ion (representing the isotope containing 1 ^13^C atom) was found in the extracts of six strains. ^28^ Again, its MS^2^ fragmentation spectrum matched the one reported in the literature (see Figures S4 and S32). Further, the thiopeptide siomycin A ([M+H]^+^ 1648.46 *m/z*), along with its congeners, siomycin B ([M+H]^+^ 1510.42 *m/z*), siomycin C ([M+2H]^2+^ 832.73 *m/z*) and siomycin D1 ([M+2H]^2+^ 817.73 *m/z*) was detected in extracts of up to 26 strains and will be described in detail below. Finally, a signal with *m/z* 303.18, corresponding to the [M+2H]^2+^ ion of ureylene-containing oligopeptide antipain, was detected in extracts of 11 strains and annotated by comparison to an authentic standard (see Figures S4 and S20). Several more signals belonging to ureylene-containing oligopeptides were identified: the antipain-like molecule KF 77AG6 ([M+H]^+^ 366.18 *m/z*, Figures S4 and S29) as well as chymostatin A/C ([M+H]^+^ 608.31 *m/z*, Figures S4 and S24) along with its congeners chymostatin B ([M+H]^+^ 594.30 *m/z*, Figures S4 and S25), chymostatinol A ([M+H]^+^ 596.32 *m/z*, Figures S4 and S26), chymostatinol B ([M+H]^+^ 610.33 *m/z*, Figures S4 and S27) and GE-20372 A/B ([M+H]^+^ 612.31 *m/z*, Figures S4 and S28). Except for thiostrepton, all previously reported *Planomonospora* metabolites were identified in our dataset. In addition, we were able to detect several members of the desferrioxamine family, an iron chelating siderophore commonly produced by *Streptomyces*.^41^ The signal corresponding to desferrioxamine B [M+H]^+^ ion (561.36 *m/z*), detected in extracts of 8 strains, was annotated by comparison to a commercial standard (see Figures S4 and S30). Further, signals matching Acyl Desferrioxamin C13, C15 and C16 ([M+H]^+^ 729.54 *m/z*, [M+H]^+^ 757.58 *m/z* and [M+H]^+^ 771.59 *m/z*, Figures S4 and S21-23, respectively) were identified by literature search. Some other putative siderophores were identified by spectral matching against the GNPS spectral library (see Figure S4): desferrioxamine E ([M+H]^+^ 601.36 *m/z)*, as well as partially described metabolites deposited as “amphiphilic ferrioxamine 7” ([M+H]^+^ 768.44 *m/z)*, “Bisu-05” ([M+H]^+^ 345.35 *m/z)* and “Desf-05” ([M+H]^+^ 575.37 *m/z*).

Several of the de-replicated features showed strong phylogroup-specificity: the desferrioxamines were almost exclusively detected in samples from strains of phylogroup “C” (see Figure 6A), while siomycins and sphaericin were only detected in extracts derived from the “S” phylogroup strains (Figure 6C). Other metabolites were less phylogroup-specific: chymostatinol A was produced by 31 strains from phylogroups “C”, “S”, “V2” and “A2”, as were antipain (11 strains, phylogroups “S”, “V2” and “A2”) or GE-20372 A/B (17 strains, phylogroups “C”, “S”, “V2” and “A2”). The other identified ureylene-containing oligopeptides showed a similar broad distribution.

In total, 28 metabolites were annotated as CAWG classes 1 or 2 (see Figure S4). A summary of the presence/absence of all de-replicated features can be seen in Figure S5. Many more features in the molecular network are neighbours to annotated ones (and thus, structurally related). While a systematic investigation of all features would exceed the scope of this study, examples of special relevance will be discussed below. To get a better picture of the biosynthetic capacities of *Planomonospora* and to further explore the annotated metabolites, we turned to genomic analysis.

### Genome analysis

Public databases report only a single *Planomonospora* genome sequence, namely that of *Planomonospora sphaerica* JCM9374. ^42^ Thus, we selected three representative strains for full genome sequencing: strain ID67723, as a representative of the divergent phylogroup “V2” and for its capability to produce chymostatin; strain ID82291 as the representative of a subgroup of strains in phylogroup “A2” that produced a family of modified peptides, as reported elsewhere;^43^ and strain ID91781, with its 16S rRNA sequence identical to that of *P. sphaerica* JCM9374 and a producer of the thiopeptide siomycin. Genomes were sequenced with both Illumina HiSeq and PacBio technologies to allow hybrid assembly, providing good quality sequences with a substantially lower number of contigs that the reference genomes of *P. sphaerica* JCM9374 and of *Planobispora rosea* ATCC53733. ^44^ The three genomes were similar to that of *P. sphaerica* JCM9374 and to each other in terms of GC content (from 71.5 to 72.83%, see Figure 7B). Interestingly, the genome of strain ID82291 was about 9% smaller than the other *Planomonospora* genomes and thus harboured a smaller number of predicted genes. antiSMASH analysis identified between 23 and 28 biosynthetic gene clusters (BGCs) in the three genomes (see Figure 7A).

**Figure 7.**
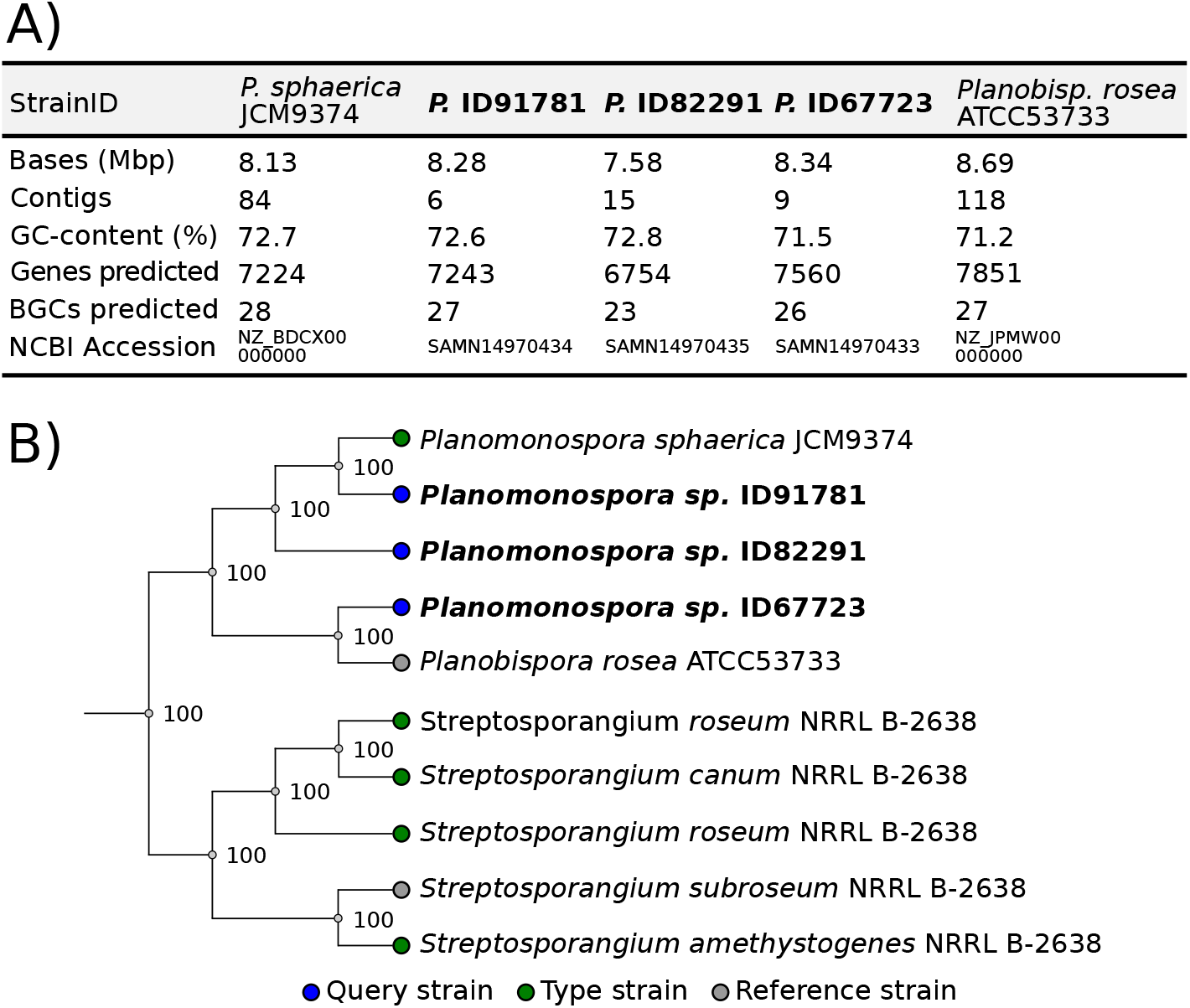
Overview of sequenced genomes, with comparison to reported ones. (A) Summary of metadata of newly sequenced genomes (bold type, P=*Planomonospora*). (B) Segment of autoMLST-generated phylogenetic tree.

A multilocus sequence analysis of the five strains of Figure 7A, along with other publicly available genome sequences of members of the *Streptosporangiaceae*, was performed with the web-based program autoMLST. ^45^ Based on a concatenated alignment of 89 identified housekeeping genes (see Figure S38), a tree was constructed that showed consistent with the one of Figure 2, except that strain ID91781 is now clearly distinct from *Planomonospora sphaerica* JCM9374 (see Figure 7B). Oddly, strain ID67723 shows closer relationship to *Planobispora rosea* than to the other *Planomonospora* strains, in contrast to the results from the 16S rRNA-based phylogeny (see Figure 2). These results warrant further studies in the taxonomy of *Planomonospora* and *Planobispora* ssp.

To investigate the similarity among the *Planomonospora* BGCs, we processed the output of antiSMASH v5.0.0^8^ with the program BiG-SCAPE/CORASON v1.0.^46^ This tool calculates sequence similarity networks and groups related BGCs in gene cluster families (GCFs), based on Pfam composition similarity. It further allows comparison against the MIBiG repository of experimentally established BGCs and automated calculations of phylogenetic relationships between BGCs.

The 131 BGCs from the genomes of Figure 7A could be grouped into 49 GCFs, of which 17 were singletons (only one member), as illustrated in Figure 8. Interestingly, the analysis demonstrated the existence of eight GCFs that are common among the four *Planomonospora* strains as well as *Planobispora rosea* ATCC53733. An additional GCF is present in all strains, except for ID82291, the strain with a slightly reduced genome. On the contrary, only two GCFs are present in the four *Planomonospora* genomes but not in *Planobispora rosea*. Only two of these eight core GCFs are highly related to experimentally established BGCs, namely those for the lantipeptide catenulipeptin and for the polyketide alkylresorcinol. Some of these GCFs are highly conserved in other genomes of members of the Streptosporangiaceae (Figure 8).

**Figure 8.**
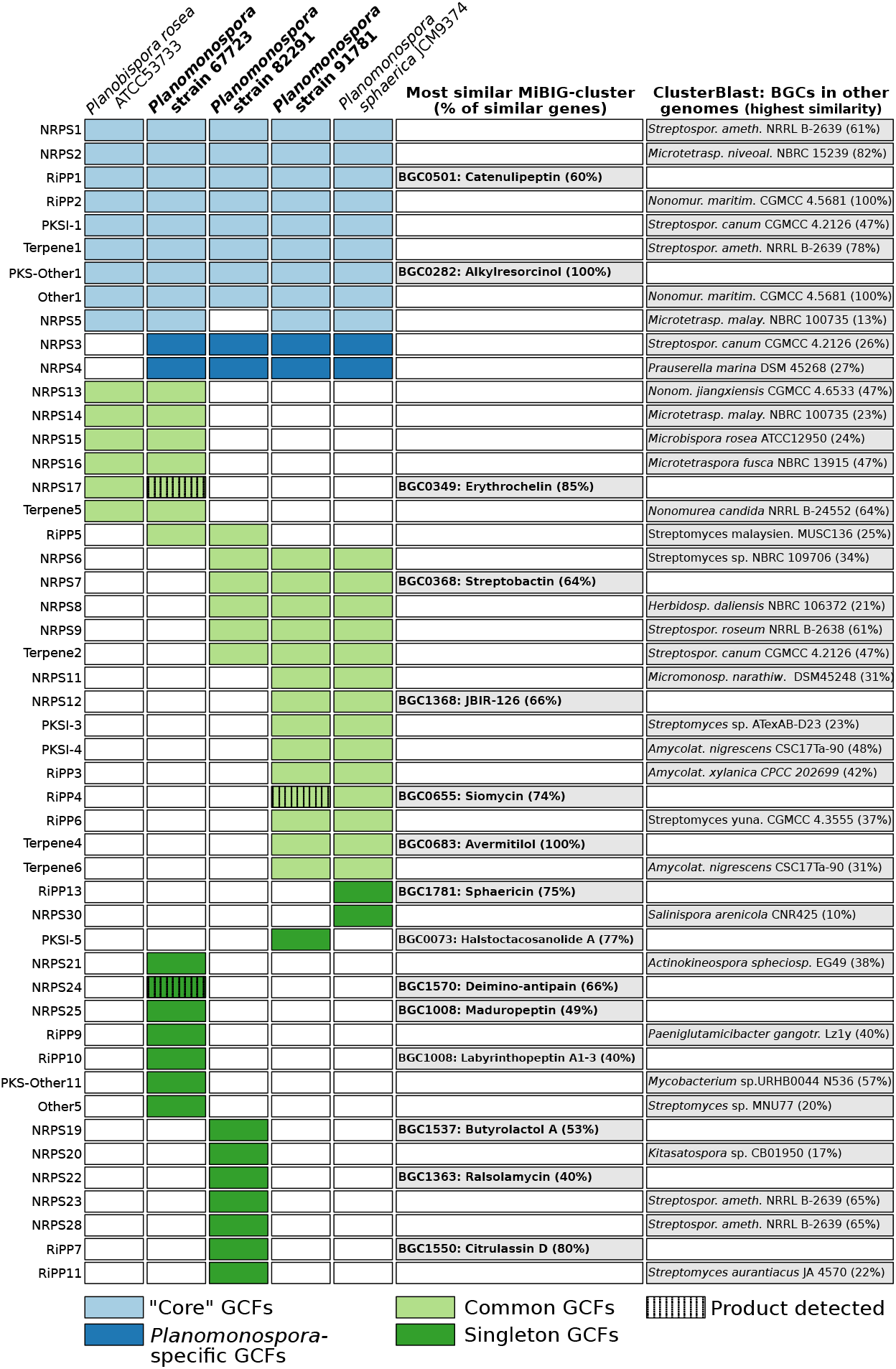
Distribution of biosynthetic gene clusters (BGCs) in the genomes of four *Planomonospora* and one *Planobispora* strains. Newly sequenced strains are indicated in bold. The comparison is based on similarity between genes and gene clusters, calculated by the program BiG-SCAPE. BGCs with a similarity of more than 40% to a MIBiG-deposited gene cluster were annotated. For clusters without a MIBiG-annotation, the next most similar BGC is indicated (found in public genomes by ClusterBlast). The twelve *Planobispora rosea* singleton BGCs were omitted, as were one BGC of strain ID67723, one of *P.sphaerica* and one of ID91781, fragmented due to their position on contig edges.

In addition to the core GCFs, strain ID67723 and *Planobispora rosea* ATCC53733 share six additional BGCs, consistent with the close relationship indicated in the autoMLST-generated phylogenetic tree. This overlap includes an erythrochelin-like BGC, as discussed later. Further, strain ID91781 and *Planomonospora sphaerica* JCM9374 are remarkably similar in terms of GCFs: ID91781 lacks the sphaericin BGC and another cluster of unknown function, present in *Planomonospora sphaerica* JCM9374, but contains instead a type I PKS BGC.

When compared to the MIBiG-repository, only 15 (30%) of all GCFs have a matching cluster. Investigation with the program ClusterBlast showed related BGCs in non-*Planomonospora* genomes. Similarities were low in the majority of cases, except for some of the core GCFs (see Figure 8).

In summary, the three analyzed genomes contain a considerable number of BGCs, with a core of BGCs present in several representatives of the Streptosporangiaceae family. Strain ID67723 appears to be very similar to *Planobispora rosea* ATCC53733, also in terms of BGCs. However, a larger number of genomes is needed to better understand the distribution of GCFs in *Planomonospora*, as demonstrated in studies on *Planctomycetes*^47^ or *Salinispora*. ^15,16^ Many of the BGCs in *Planomonospora* remain unannotated and may encode for novel metabolites. Aiming for a better understanding of the biosynthetic capabilities, we connected genomic and metabolomic data creating a “paired-omics” dataset, as illustrated below.

### Paired -omics

Siomycin, first reported from *Planomonospora* in 1968 as sporangiomycin,^23^ is a thiostrepton-like thiopeptide with an established biosynthetic route.^48^ While the siomycin BGC was detected in strain ID91781 (RiPP4 in Figure 8), during metabolite annotation, it became evident that the features matching siomycin A and congeners B, C and D1 were not clustered in a single molecular family in the molecular network, as expected for structurally related molecules. Instead, they were mostly singletons. Inspection of the corresponding MS^2^ spectra revealed that siomycin A and congeners shared low molecular weight fragments, probably corresponding to the quinalidic acid (QA) moiety of class b thiopeptides (see Figure S6). Hence, the program MS2LDA^49^ was used to mine for QA-related motifs in the MS^2^ fragmentation spectra of features. Apart from the known siomycins, the program detected further nine putative siomycin-like thiopeptides (see Figure S7). One of them, which we named siomycin E, is hypothesized to correspond to siomycin B with one additional dehydroalanine (Dha)-residue at its C-terminal end, instead of two, as in siomycin A. MS^2^-fragmentation spectra of the [M+H]^+^ ions showed a particular fragment corresponding to a break between Ala^2^- and Dha^3^ as well as Thr^12^ and QA, with an *m/z*-value diagnostic for each congener (see Figures 9 and S8). Literature provides a precedent for a thiopeptide with a similar intermediate molecule: thiopeptin A3a, A4a and Ba have zero, one and two Dha residues, respectively, at the C-terminal end of the molecule.^50^ The precursor peptide for thiopeptin, TpnA, indeed shows two serine moieties at the C-terminal end. ^51^ Consistently, the precursor peptide encoded by the BGC RiPP4 contains two additional serine residues at the C-terminus (see Figure S9). Promiscuous processing of the C-terminal end of precursor peptides has been observed in several thiopeptides and can be also assumed here. ^48,50,52,53^ For some of the other putative thiopeptides, the differences in exact mass with regard to siomycin A point towards one or two *N*-acetylcysteinyl-moieties with additional modifications, such as deacetylation or hydroxylation (see Figure S7). Again, literature provides precedence for likewise modified antibiotics from *Streptomyces* sp.: *N*-acetylcysteine derivates have been isolated for the macrolide piceamycin^54^ and the phenazine SB 212021.^55^ These results show how the use of additional tools such as MS2LDA can overcome the limitations of any one tool and help to organize data, identify analogues and support annotation (for a full list of annotated Mass2Motifs, see Figure S10).

**Figure 9.**
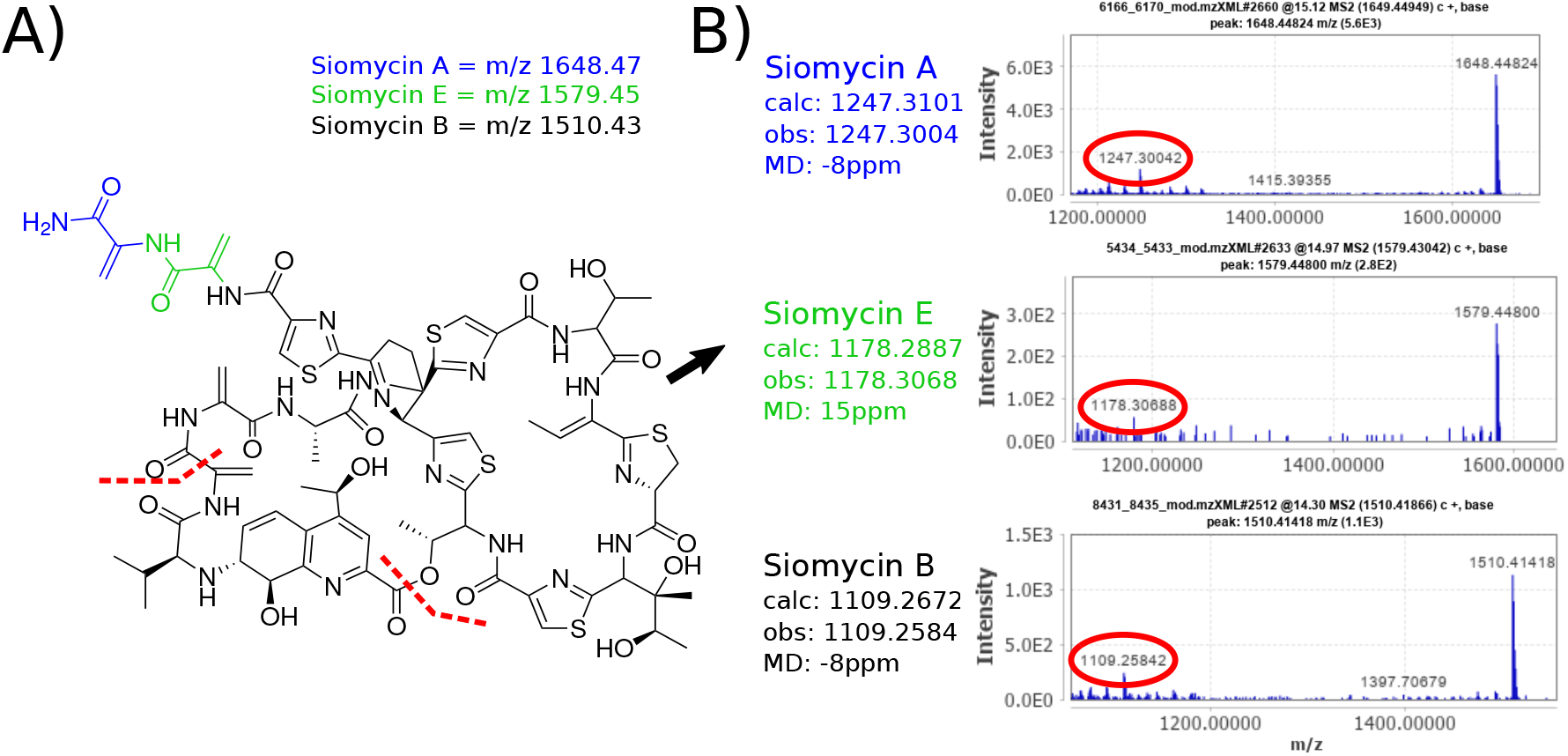
Identification and annotation of siomycin congeners. (A) shows the putative tandem mass fragmentation pathway of siomycins, leading to the diagnostic fragments indicated in (B). Siomycin A (blue) has two dehydroalanine (Dha)-moieties at its C-terminus end, while siomycin B (black) has none. Siomycin E (green) is hypothesized to be an intermediate congener with one Dha-group at its C-terminal end.

As mentioned above, several features were found to match desferrioxamines, iron-chelating siderophores involved in iron-uptake in bacteria,^56,57^ mostly in samples derived from strains of phylogroup “C”, raising the question if the other *Planomonospora* strains produced distinctive iron-chelating molecules. In a study on 118 *Salinispora* genomes, Bruns et al. reported a mutually exclusive presence of either the *des* or *slc* BGC, responsible for the production of desferrioxamine or the structurally unrelated salinichelins, respectively. ^58^ In order to rapidly identify iron-binding metabolites, we added FeCl3 to the *Planomonospora* extracts and reanalyzed them by LC-MS/MS, looking for mass shifts from disappearance of the iron-free form and stabilization of the iron-bound molecule, with associated Fe-characteristic isotopic pattern. Apart from the already identified desferrioxamines, we found that 18 features of four molecular families were also affected by the addition of iron. (see Figure 10A and Figures S12 and S14-S19). Calculation of molecular formulas and inspection of MS^2^-fragmentation spectra indicated relatedness between these four molecular families, but not to desferrioxamine E (see Figures S12 and S13). These four families were exclusive to 7 strains belonging to phylogroups “V2” and “S”. At the same time, no desferrioxamines could be detected in extracts from these strains. Analysis with BiG-SCAPE/CORASON suggested a candidate BGC (NRPS17 in Figure 8) in one of the producer strains, ID67723. This cluster showed high similarity to the experimentally validated BGC for erythrochelin, a siderophore produced by *Saccharopolyspora erythraea*, as well as to a BGC with unknown function from *Planobispora rosea* ATCC53733, as indicated in Figure 10B. Consistently, extracts from *Planobispora rosea* also contained a feature with *m/z* 631.3408 [M+H]^+^ with the calculated molecular formula C_26_H_46_N_8_O_10_ (see Figure 10A). Characterisation of this metabolite confirmed a salinichelin-like structure, with a lysine instead of an arginine in position two (manuscript in preparation). For strain ID67723, 11 additional iron-shifted features were identified. The similarity of their MS^2^ fragmentation spectra to the features with *m/z* 631.3408 [M+H]^+^ suggests that they were also synthesized by BGC NRPS17. While many were hypothesized to be congeners differing in one or more methylene groups, feature ID314 (*m/z* 1067.5659) appears to be a glycosylated version of feature ID289 (*m/z* 905.5136).

**Figure 10.**
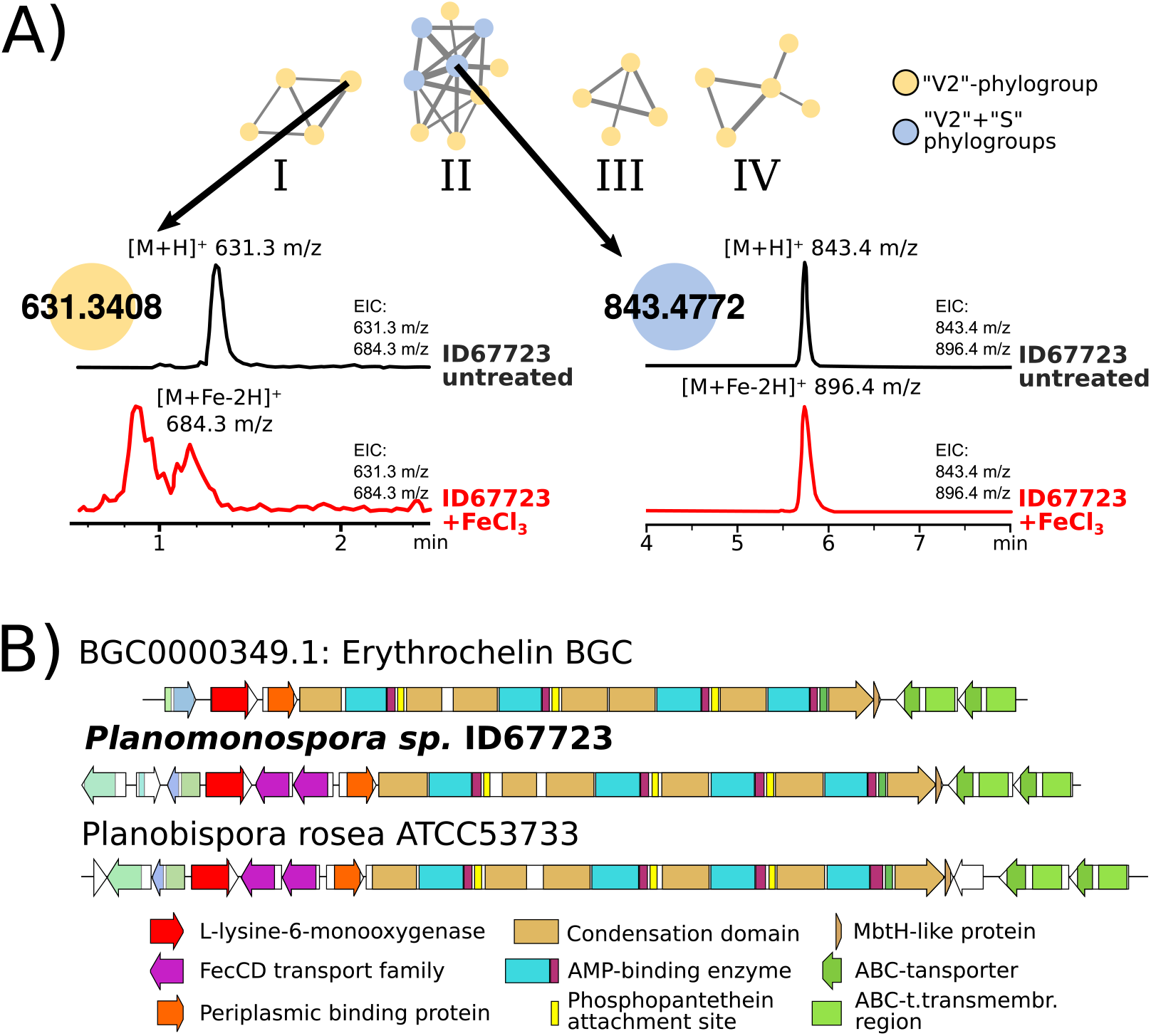
Investigation of some iron-binding metabolites: (A) features in several molecular families, such as *m/z* 631.3 or *m/z* 843.4, showed iron complexion upon treatment with FeCl_3_. LCMS-analysis of Fe-treated samples show disappearance of the unbound form (black trace) and appearance of the Fe-bound form (red trace; for original traces, see Figure S11). (B) Comparison of the erythrochelin BGC with GCF NRPS17, shared by *Planomonospora sp*. ID67723 and *Planobispora rosea* ATCC53733.

## Conclusion

The work presented here allowed us to get unprecedented insight into the poorly explored genus *Planomonospora*. Among the 72 investigated strains, we found phylogenetic diversity that exceeds the (few) known isolates. Using feature-based molecular networking, we could find the majority of previously reported *Planomonospora* metabolites, as well as demonstrating phylogroup-specificity for many of them. We also showed that *Planomonospora* can produce desferrioxamines as iron-chelating molecules, or, alternatively, members of a different siderophore family. The majority of features detected in the *Planomonospora* extracts and of the BGCs of the available *Planomonospora* genomes remain unknown, underlying the very limited knowledge we have about this genus.

## General Experimental Procedures

### Cultivation

Strains from frozen stocks (−80°C) were propagated on S1 plates^59^ at 28°C for two to three weeks. The grown mycelium was then homogenized with a sterile pestle and used to inoculate 15 mL AF (=AF/MS)^59^ medium in a 50 mL baffled flask. After cultivation on a rotary shaker (200 rpm) at 30°C for 72 hours, 1.5 ml of the exponentially growing culture was used to inoculate each 15 mL of MC1 (35 g/L soluble starch, 10 g/L glucose, 2 g/L hydrolyzed casein, 3.5 g/L meat extract, 20 g/L yeast extract, 10 g/L soybean meal, 2 g/L CaCO_3_, adjusted to pH 7.2), AF2 (8g/L yeast extract, 30 g/L soybean meal, 11 g/L glucose, 25 g/L malt extract, 1 g/L L-valine, 0.5g/L Biospumex (Cognis, France), adjusted to pH 7.4), R3 (=RARE3)^60^ and AF media in a 50 mL baffled flask. After 7 days of cultivation as before, cultures were harvested and extracted as described below.

### Extraction and LC-MS sample preparation

10 mL of each culture was centrifuged at 16,000 rcf for 10 min and the resulting pellet separated from the supernatant. For each culture, two different extracts were prepared: one by solvent extraction of the mycelium, and one by solid-phase adsorption of the cleared broth. The mycelium was resuspended in 4 mL EtOH and incubated under agitation at room temperature for one hour. After centrifugation (16.000 rcf for 10 min), the pellet was discarded and the mycelium extract was processed as described below. The cleared broth was stirred with 1 mL of HP20 resin suspension (Diaion) for one hour at room temperature. Then, the exhausted broth was discarded and the loaded HP20 resin was stirred with 4 mL of purified water (MilliQ, Merck), then recovered by centrifugation and eluted with 5 mL 80:20 MeOH:H_2_O under agitation at room temperature for 10 min. Both extracts were transferred to 96-well plates (conic bottom). For mycelium extract, 100 μL of extract were deposited, while for supernatant extract, 125 μL were deposited. Extracts were dried under vacuum at 40°C. Before analysis, extracts were rehydrated: for mycelium-extracts, 100 μL 90% EtOH was used, while for supernatant-extracts, 125 μL 80% MeOH was utilized. The samples were centrifuged for 3 min at 16,000 rcf to remove suspended particles.For LCMS-analysis, 50 μL of mycelium extract and 50 μL of supernatant extract, coming from a single culture, were combined, centrifuged for 3 min at 16.000 rcf and transferred to HPLC vials for analysis. To test for iron-binding metabolites, 10 μL 10 mg/mL FeCl3 in water (MilliQ, Merck) was added to selected rehydrated extracts.

### LC-MS/MS analysis

For the 286 *Planomonospora* extracts investigated in molecular networking, data acquisition was performed on a Dionex UltiMate 3000 (Thermo Scientific) coupled with a micrOTOF-Q III (Bruker) instrument equipped with an electrospray interface (ESI). Samples were analyzed on an Atlantis T3 C18 5μm x 4.6 mm x 50 mm column maintained at 25°C with a flow rate of 0.3mL/min. Phases A and B were 0.1% acetic acid in water and 0.1% acetic acid in acetonitrile (Sigma), respectively. All solvents used were LCMS-grade. A multistep program was followed that consisted of 10, 10, 100, 100, 10, 10 % phase B at 0, 1, 20, 26, 27, 30 min, respectively. MS data were acquired in data dependent acquisition (DDA) mode over a range from 100-3000 *m/z* in positive ionization mode. Auto MS/MS fragmentation was achieved with rising collision energy (35-50 keV over a gradient from 500-2000 *m/z*) with a frequency of 4 Hz for all ions over a threshold of 100. For the analysis of commercial standards and selected *Planomonospora* extracts with added FeCl3, LC-MS analyses were performed on a Dionex UltiMate 3000 (Thermo Scientific) coupled with an LCQ Fleet (Thermo scientific) mass spectrometer equipped with an electrospray interface (ESI) on an Atlantis T3 C_18_ 5μm x 4.6 mm x 50 mm column, as reported elsewhere.^61^

### HR-LC-MS/MS data calibration and conversion

LCMS/MS-files, coming from the high resolution micrOTOF-Q III (Bruker) instrument, were calibrated with Bruker Data-Analysis using an internal calibrant (Na-acetate) in HPC mode, injected at the beginning of each run. The calibration was verified by inspection of medium component soyasaponin A, present in all samples (calc. [M+H]^+^ 943.526 *m/z* at RT 12.7-12.9min). Average mass deviation was routinely below 15ppm. LCMS files were exported (see Figure S35) to the .mgf and .mzXML format with Bruker DataAnalysis (version 4.2 SR2 Build 365 64bit) and further processed using an *ad hoc*-written Perl5 script (see Figure S35 and S36). This was necessary for further processing with MZmine2, since the export to the .mzXML format erroneously resulted in the insertion of the non-calibrated value in the precursor-entry (`<precursorMz></precursorMz>`) of each MS^2^-scan, leading to high mass deviations (>100ppm) and disrupting the logical connection between MS^1^ and MS^2^ scans in the .mzXML files. .mgf files were not affected by this and thus, our script replaces the non-calibrated entries in the .mzXML file with the correct ones from the .mgf file.

### Pre-processing of HR-LC-MS/MS data with MzMine2

For pre-processing with MZmine2, mzXML files were imported and subjected to the following workflow: A) Mass-Detection: Retention time: auto; MS^1^ noise level: 1E3; MS^2^ noise level: 2E1. B) ADAP chromatogram builder:^62^ Retention time: auto; MS-level: 1; Min group size in # of scans: 8; Group intensity threshold: 5E2; Min highest intensity: 1E3; *m/z* tolerance: 20ppm; C) Chromatogram deconvolution: Baseline-cutoff-algorithm; Min peak height: 1E3; Peak duration: 0.1-1.3 min; Baseline level: 2.5E2; *m/z* range MS^2^ pairing: 0.02; RT range MS^2^ pairing: 0.4 min; D) Isotopic peaks grouper: *m/z* tolerance: 20ppm; RT tolerance: 0.2 min; Monotonic shape: no; Maximum charge: 2; Representative isotope: Most intense; E) RANSAC peak alignment: *m/z* tolerance: 20 ppm; RT tolerance: 0.7 min; RT tolerance after correction: 0.35 min; RANSAC iterations: 100000; Minimum number of points: 50%; Threshold value: 0.5; Linear model: no; Require same charge state: no; F) Duplicate peak filter: Filter mode: New average; *m/z* tolerance: 0.02 *m/z*; RT tolerance: 0.4 min. Features with no accompanying MS^2^ data were excluded from the analysis. Features present in both samples and media blanks were excluded as well, since they related to media components. Features with *m/z*-values of <300 or a retention time <1.5 min or >20 min were also excluded. The resulting feature list contained 1492 entries and was exported to the GNPS-compatible format.

### Global Natural Products Social Molecular Networking (GNPS) feature-based molecular MS/MS network

Using the Feature-Based Molecular Networking (FBMN) workflow (version release_14)^35^ on GNPS,^12^ a molecular network was created by processing the output of MZmine2. Parameters were adapted from the GNPS documentation: MS^2^ spectra were filtered so that all MS/MS fragment ions within +/− 17 Da of the precursor *m/z* were removed and only the top 6 fragment ions in the +/− 50 Da window through the spectrum were utilized, with a minimum fragment ions intensity of 50. Both the MS/MS fragment ion tolerance and the precursor ion mass tolerance were set to 0.03 Da. Edges of the created molecular network were filtered to have a cosine score above 0.7 and more that 5 matched peaks between the connected nodes. Further, these were only kept in the network if each of the nodes appeared in each other’s respective top 10 most similar nodes. The maximum size of molecular families in the network was set to 250 and the lowest scoring edges from each family were removed until member count was below this threshold. The MS^2^ spectra in the molecular network were searched against GNPS spectral libraries. ^12,63^ Reported matches between network and library spectra were required to have a score above 0.6 and at least 5 matched peaks. The DEREPLICATOR-program was used to annotate MS/MS spectra.^64^ The molecular networks were visualized using Cytoscape 3.7.1.^36^ The molecular networking job is accessible by the link https://gnps.ucsd.edu/ProteoSAFe/status.jsp?task=92036537c21b44c29e509291e53f6382. HR-ESI-LC-MS/MS data were deposited in MassIVE (MSV000085376) and linked with the genomic data at the iOMEGA Pairing Omics Data Platform (http://pairedomicsdata.bioinformatics.nl/projects).

### MS2LDA analysis

The molecular networking job described above was analyzed by MS2LDA (version release_14), accessing the tool directly on the GNPS website. Parameters were set as follows: Bin Width: 0.01; Nr of LDA iterations: 1000; Min MS2 Intensity: 100; LDA Free Motifs: 500. All MotifDBs except “Streptomyces and Salinisporus Motif Inclusion” were excluded. Further parameters were left at default (Overlap score threshold: 0.3; Probability value threshold: 0.1; TopX in node: 5). Results were uploaded to the MS2LDA-website, with the width of ms2 bins set to 0.005 Da, as recommended (http://ms2lda.org/).

### 16S rRNA gene analysis

For 16S rRNA gene amplification, single colonies were picked from S1 medium plates and lysed at 95°C in 100 μL PCR-grade water for 5 min. 5 μL centrifuged lysate was added to the reaction mix, containing 25 μL DreamTaq Green PCR Master Mix 2X (Thermo Scientific), 3 μL of 10 x Denhardt’s reagents,^65^ each 500 nM of eubacterial primers R1492 and F27^66^ and 12 μL water, resulting in a final volume of 50 μL. The amplification was performed as reported elsewhere. ^66^ PCR products were sequenced using Sanger sequencing by an external DNA sequencing provider (Cogentech, Milan, IT), with the primers mentioned above. 16S rRNA gene sequences were inspected and assembled manually using the software AliView, ^67^ yielding 35 non-redundant sequences with a consensus length of 1377 bp. The sequences were analyzed with programs contained in the PHYLIP package^68,69^ as reported elsewhere,^66^ with slight modifications (bootstrapped with 1000 replicates). The resulting consensus tree was visualized using the iTOL-webserver.^70,71^ Sequences were deposited in GeneBank, with accession numbers included in the supplemental information (Figures S1-2) of this study.

### Isolation of gDNA

To isolate genomic DNA (gDNA), mycelium from 5 mL *Planomonospora* cultures, cultivated in AF medium for 72 hours as described above, was extracted with standard protocols for *Streptomyces* by phenol-chloroform, as described elsewhere. ^44^ gDNA was sequenced with both Illumina and PacBio technologies by an external service provider (Macrogen, Seoul, KOR) and assembled using the program SPAdes (3.11). Genome sequences were deposited in GenBank under the BioProject ID PRJNA633779, with accession numbers JABTEX000000000 (ID82291), JABTEY000000000 (ID91781) and JABTEZ000000000 (ID67723).

### BiG-SCAPE sequence similarity network

.gbk-files of BGCs detected by the webbased application antiSMASH 5.0 (https://antismash.secondarymetabolites.org/) were analyzed including the ClusterFinder algorithm and compared to MIBiG-deposited BGCs using BiG-SCAPE 1.0 in “hybrids” mode. All setting were left to default, except the cutoff-value, which was set to 0.5.

### Phylogenetic multilocus sequence analysis (autoMLST)

The web-based application autoMLST (http://automlst.ziemertlab.com/analyze#) was used to perform a multilocus sequence analysis in “denovo mode” and default settings.

## ACKNOWLEDGEMENT

The authors thank Christian Milani and Marco Ventura for genome reassembly and data curation, Paolo Monciardini for suggestions regarding the phylogenetic analysis and the whole Naicons team for helpful comments and discussion. M.M.Z. thanks Prof. Tanneke den Blaauwen for guidance and advice. This work has received funding from the European Union’s Horizon 2020 research and innovation program under grant agreement No.721484 (Train2Target). M.C. received funding from the German Research Foundation (DFG) grant nr. FOR2372.

## ASSOCIATED CONTENT

The following files are available free of charge.

- Supporting Information: Figures S1-S38.

## Notes

### Competing Interest Statement

The authors have declared no competing interest.

