## Supporting Information for "*Planomonospora*: a Metabolomics Perspective on an Underexplored Actinobacteria Genus"

### *Planomonospora*: a Metabolomics Perspective on an Underexplored Actinobacteria Genus - Supporting Information

#### Strain-list 1/2

| Strain ID | Origin | Source[a] | Reception[b] (Y-M-D) | Phylo-group[c] | Most similar strain[d] (%) | NCBI Access.[e] |
|---|---|---|---|---|---|---|
| 103354 | Costa Rica | Soil, Coco, Nicoya Peninsula | 1996-07-17 | A2 | *P.algeriensis* PM3 (99.55) | MT375135 |
| 103452 | Niger | Soil, millet field, Boubon | 1996-08-08 | A1 | *P.sphaerica* JCM9374 (99.48) | MT375136 |
| 103466 | Niger | Soil, millet field, Boubon | 1996-08-08 | A1 | ID103452 (100) | |
| 107089 | Italy | Soil, vineyard, Siena | 1996-08-18 | A2 | *P.algeriensis* PM3 (99.63) | MT375137 |
| 107144 | Niger | Soil, Tamaské | 1996-08-08 | A2 | *P.algeriensis* PM3 (99.70) | MT375138 |
| 107147 | Niger | Soil, Tamaské | 1996-08-08 | A2 | ID98083 (100) | |
| 107158 | Niger | Soil, Tamaské | 1996-08-08 | C | *P.corallina* A-T 11038 (99.93) | MT375139 |
| 107159 | Niger | Soil, Tamaské | 1996-08-08 | C | *P.corallina* A-T 11038 (99.85) | MT375140 |
| 107162 | Niger | Soil, Tamaské | 1996-08-08 | A2 | ID98083 (100) | |
| 107188 | Tanzania | Soil, Lake Manyara | 1997-08-01 | S | ID91781 (100) | |
| 108730 | India | Soil, rice field | 1998-08-31 | S | *P.sphaerica* JCM9374 (99.78) | MT375141 |
| 1089061 | Israel | Soil, Golan Heights | 1998-08-31 | S | ID1089062 (100) | |
| 1089062 | Israel | Soil, Golan Heights | 1998-08-31 | S | *P.sphaerica* JCM9374 (99.93) | MT375142 |
| 114239 | India | Soil, rice field | 1998-08-31 | A2 | *P.algeriensis* PM3 (99.55) | MT375143 |
| 114465 | Morocco | Soil, Zagora | 1998-08-31 | V2 | ID67723 (100) | |
| 114499 | Malta | Soil, Mdina | 1998-08-31 | A2 | *P.algeriensis* PM3 (99.93) | MT375169 |
| 119016 | Unknown | INA collection Moscow | 1995-03-14 | S | ID91781 (100) | |
| 119052 | Unknown | INA collection Moscow | 1995-03-14 | C | *P.corallina* A-T 11038 (99.63) | MT375144 |
| 1299671 | Italy | Soil, Matera | 2001-03-01 | V2 | ID67723 (100) | |
| 1299672 | Italy | Soil, Matera | 2001-03-01 | V2 | ID67723 (100) | |
| 131892 | Italy | Soil, Langhe, Asti | 2001-04-19 | S | *P.sphaerica* JCM9374 (99.85) | MT375145 |
| 131917 | Egypt | Soil, Hurghada, Egypt | 2000-09-07 | S | *P.sphaerica* JCM9374 (99.85) | MT375146 |
| 134022 | India | Soil | 1978-03-01 | S | ID91781 (100) | |
| 134035 | India | Soil | 1978-03-01 | S | ID91781 (100) | |
| 134049 | India | Soil | 1978-03-01 | A2 | *P.algeriensis* PM3 (99.78) | MT375147 |
| 134050 | India | Soil | 1978-03-01 | S | ID91781 (100) | |
| 135044 | Brazil | Soil, Amazonas | 1994-09-26 | C | *P.corallina* A-T 11038 (99.70) | MT375148 |
| 135062 | Nicaragua | Soil, Salamasi | 1999-05-07 | S | ID91781 (100) | |
| 43178 | Unknown | Lepetit Historic Collection | Unknown | V1 | *P.venezuelensis* JCM 3167 (99.93) | MT375149 |
| 43517 | Unknown | Lepetit Historic Collection | Unknown | S | ID88009 (100) | |
| 43657 | Unknown | Lepetit Historic Collection | Unknown | S | *P.sphaerica* JCM9374 (99.85) | MT375150 |
| 46103 | Unknown | Lepetit Historic Collection | Unknown | S | *P.sphaerica* JCM9374 (99.85) | MT375151 |
| 46108 | Unknown | Lepetit Historic Collection | Unknown | S | *P.sphaerica* JCM9374 (99.78) | MT375152 |
| 46114 | Unknown | Lepetit Historic Collection | Unknown | S | ID91781 (100) | |
| 46116 | Unknown | Lepetit Historic Collection | Unknown | A2 | ID82291 (100) | |
| 50035 | Unknown | Lepetit Historic Collection | Unknown | S | ID91781 (100) | |

[a]Source and place of sampling, if known, OR strain collection, originating from.
[b]Date of addition to strain library.
[c]Taxonomic group used in this publication.
[d]Most similar sequence (type strain or representative strain, described in this study), with % of sequence identity indicated.
[e]Accession number of deposited sequence. Only representative, non-redundant sequences were deposited.

Figure 1: List of strains investigated in this study.

#### Strain-list 2/2

| Strain ID | Origin | Source[a] | Reception[b] (Y-M-D) | Phylo-group[c] | Most similar strain[d] (%) | NCBI Access.[e] |
|---|---|---|---|---|---|---|
| 50037 | Unknown | Lepetit Historic Collection | Unknown | C | *P.alba* JCM9373 (100) | MT375153 |
| 67723 | Greece | Soil, chard field | 1992-08-12 | V2 | *P.venezuelensis* JCM 3167 (98.81) | MT375154 |
| 69121 | Italy | Soil, wheat field, Lecce | 1992-08-18 | S | ID88035 (100) | |
| 71637 | Kenya | Soil | 1992-08-23 | S | ID91781 (100) | |
| 73480 | Italy | Soil, pine forest, Lecce | 1992-08-30 | S | ID88035 (100) | |
| 73756 | Albania | Soil, pine forest | 1993-08-12 | S | ID88009 (100) | |
| 73758 | Albania | Soil, pine forest | 1993-08-12 | S | ID88009 (100) | |
| 74193 | Italy | Soil, S. Maria die Leuca | 1993-08-12 | S | ID88035 (100) | |
| 74194 | Italy | Soil, S. Maria die Leuca | 1993-08-12 | S | ID88035 (100) | |
| 75183 | Pakistan | Soil, Peshawar | 1993-08-12 | S | ID91781 (100) | |
| 75185 | Pakistan | Soil, Peshawar | 1993-08-12 | C | *P.corallina* A-T 11038 (99.26) | MT375155 |
| 81182 | Unknown | INA collection Moscow | 1995-03-14 | S | *P.sphaerica* JCM9374 (99.85) | MT375156 |
| 82291 | Unknown | INA collection Moscow | 1995-03-14 | A2 | *P.algeriensis* PM3 (99.48) | MT375157 |
| 82300 | Unknown | INA collection Moscow | 1995-03-14 | C | *P.corallina* A-T 11038 (99.55) | MT375158 |
| 88009 | Italy | Soil, clover field, Cantone | 1993-08-12 | S | *P.ssp.antibiotica* JCM3094 (99.93) | MT375159 |
| 88010 | Italy | Soil, clover field, Cantone | 1993-08-12 | S | *P.sphaerica* JCM9374 (99.85) | MT375160 |
| 88035 | Nicaragua | Soil, San Juan del Norte | 1994-03-02 | S | *P.sphaerica* JCM9374 (99.78) | MT375161 |
| 91428 | Kenya | Soil, forest | 1995-03-19 | V2 | ID67723 (100) | |
| 91429 | Tanzania | Soil, Mwanza | 1995-03-19 | S | ID91781 (100) | |
| 91430 | Tanzania | Soil, Mwanza | 1995-03-19 | S | ID91781 (100) | |
| 91431 | Tanzania | Soil, Mwanza | 1995-03-19 | S | ID91781 (100) | |
| 91432 | Tanzania | Soil, Mwanza | 1995-03-19 | S | *P.sphaerica* JCM9374 (99.70) | MT375162 |
| 91433 | Tanzania | Soil, Mwanza | 1995-03-19 | S | ID91781 (100) | |
| 91525 | USA | Soil, forest, Arizona | 1995-08-20 | S | ID91781 (100) | |
| 91602 | Costa Rica | Soil, Coco, Nicoya Peninsula | 1996-07-16 | S | *P.sphaerica* JCM9374 (99.85) | MT375163 |
| 91603 | Costa Rica | Soil, Coco, Nicoya Peninsula | 1996-07-16 | S | ID91781 (100) | |
| 91611 | Costa Rica | Soil, Coco, Nicoya Peninsula | 1996-07-16 | S | *P.sphaerica* JCM9374 (99.93) | MT375164 |
| 91781 | Nicaragua | Soil | 1994-02-02 | S | *P.sphaerica* JCM9374 (100) | MT375165 |
| 96662 | Nicaragua | Soil | 1996-11-12 | C | *P.corallina* A-T 11038 (99.78) | MT375166 |
| 97500 | Brazil | Soil, Amazonas | 1995-08-13 | C | *P.corallina* A-T 11038 (99.70) | MT375167 |
| 98072 | Greece | Soil, olive garden | 1991-07-08 | V2 | ID67723 (100) | |
| 98083 | Turkey | Soil, wheat field, Izmir | 1992-05-25 | A2 | *P.algeriensis* PM3 (99.85) | MT375168 |
| 98084 | Turkey | Soil, wheat field, Izmir | 1992-05-25 | S | ID88009 (100) | |
| 98085 | Turkey | Soil, wheat field, Izmir | 1992-05-25 | S | ID88009 (100) | |
| 98216 | Syria | Soil, desert, Apamea | 1997-08-01 | S | ID88009 (100) | |
| 98217 | Syria | Soil, desert, Apamea | 1997-08-01 | S | ID88009 (100) | |

[a]Source and place of sampling, if known, OR strain collection, originating from.
[b]Date of addition to strain library.
[c]Taxonomic group used in this publication.
[d]Most similar sequence (type strain or representative strain, described in this study), with % of sequence identity indicated.
[e]Accession number of deposited sequence. Only representative, non-redundant sequences were deposited.

Figure 2: List of strains investigated in this study.

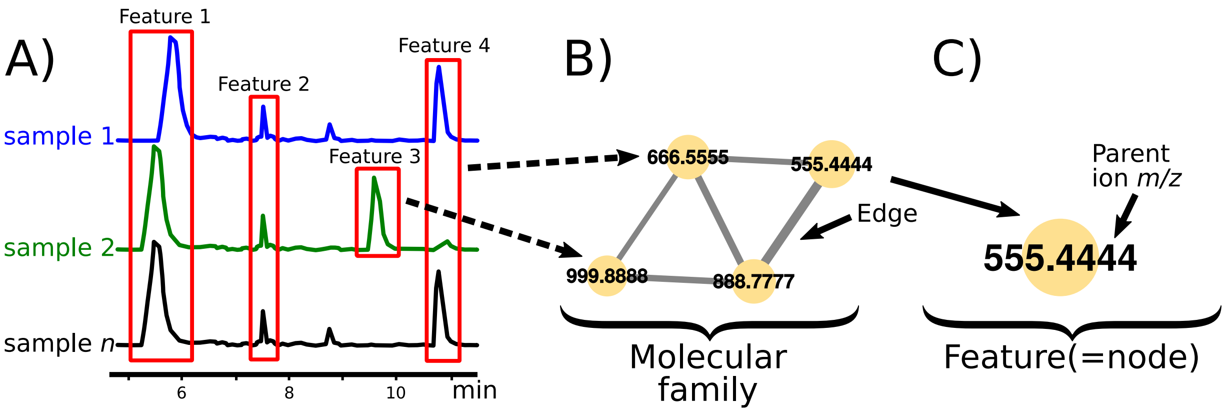

Figure 3: Schematic visualisation of feature finding and molecular networking. (A) extracted ion chromatograms from LCMS-samples are aligned by MZmine2 and peaks with common retention time and $m/z$ are detected. $MS^2$ fragmentation spectra are associated to the newly created, non-redundant features. (B) $MS^2$ fragmentation spectra of features are compared with each other and a molecular network is created, with features sporting similar $MS^2$ fragmentation pattern connected by edges. The thickness of edges correlates with the similarity between spectra. (C) Parent ion (=feature) $m/z$ is mapped onto the nodes, using the program Cytoscape.

#### Compound identification

| Feature ID[a] | Compound name (CAWG classification)[b] | Parent ion (*m/z*) | Parent ion Adduct | Reference ([M+H]+) | *m/z* error[c] | Sim. score[d] | GNPS reference[e] | Reference(s)[f] |
|---|---|---|---|---|---|---|---|---|
| 1 | Chymostatinol A (**1**) | 596.3151 | [M+H]+ | 596.315 | 0 | 0.77 | CCMSLIB00000577828 | - |
| 313 | Cholic acid[g] (2) | 817.5769 | [2M+H]+ | 409.295 | 6 | 0.94 | CCMSLIB00005465298 | - |
| 321 | Glycocholic acid (2) | 466.3137 | [M+H]+ | 466.315 | 2 | 0.74 | CCMSLIB00003139929 | - |
| 34 | Riboflavin (2) | 377.1450 | [M+H]+ | 377.145 | 0 | 0.90 | CCMSLIB00003139347 | - |
| 148 | Coproporphyrin I (2) | 328.1414 | [M+2H]2+ | 655.279 | 5 | 0.87 | CCMSLIB00003136262 | - |
| 68 | 1-Eicosatrienoyl-sn-glycerol -3-phosphoethanolamine (2) | 504.3037 | [M+H]+ | 504.306 | 4 | 0.85 | CCMSLIB00003136028 | - |
| 1070 | Desf-05 (2) | 575.3721 | [M+H]+ | 575.376 | 6 | 0.81 | CCMSLIB00003739977 | - |
| 740 | Bisu-05 (2) | 345.2486 | [M+H]+ | 345.246 | 7 | 0.69 | CCMSLIB00003739992 | - |
| 634 | Desferrioxamine E (2) | 601.3561 | [M+H]+ | 601.358 | 3 | 0.67 | CCMSLIB00000001621 | - |
| 755 | Desferrioxamine B (**1**) | 561.3574 | [M+H]+ | 561.359 | 2 | 0.64 | CCMSLIB00003134607 | - |
| 618 | Amphiphilic ferioxamine 7 (2) | 768.4403 | [M+H]+ | 768.47 | 38 | 0.6 | CCMSLIB00000077244 | - |
| 396 | Chymostatin B (**1**) | 594.2993 | [M+H]+ | 594.3034 | 7 | - | **CCMSLIB00005716838** | 10.7164/antibiotics.26.625 |
| 70 | Chymostatinol B (**1**) | 610.329 | [M+H]+ | 610.3347 | 9 | - | **CCMSLIB00005716839** | 10.7164/antibiotics.26.625 |
| 129 | Chymostatin A/C (**1**) | 608.3111 | [M+H]+ | 608.3191 | 13 | - | **CCMSLIB00005716840** | 10.7164/antibiotics.26.625 |
| 26 | GE-20372-A/B (2) | 612.3102 | [M+H]+ | 612.314 | 6 | - | **CCMSLIB00005716841** | 10.7164/antibiotics.48.332 |
| 164 | KF 77AG6 (2) | 366.1769 | [M+H]+ | 366.1772 | 1 | - | **CCMSLIB00005716842** | Watanabe Tetrahedr. 1982, 38, 1775 |
| 125 | Antipain (**1**) | 303.1788 | [M+2H]2+ | 605.3518 | 2 | - | **CCMSLIB00005716843** | 10.7164/antibiotics.25.263 |
| 1367 | 97518/Planosporicin (2) | 1097.3909 | [M+1+2H]2+ | 2192.7864 | 9 | - | **CCMSLIB00005716845** | 10.1021/np800794y |
| 16 | Riboflavol (2) | 393.1389 | [M+H]+ | 393.1403 | 4 | - | **CCMSLIB00005716846** | Pubchem CID: 129889088 |
| 1042 | Riboflavin-glucoside (2) | 539.2006 | [M+H]+ | 539.1983 | 4 | - | **CCMSLIB00005716847** | Pubchem CID: 20640019 |
| 748 | Acyl Desferrioxamin C13 (2) | 729.5448 | [M+H]+ | 729.549 | 6 | - | **CCMSLIB00005716848** | 10.1002/anie.201101225 |
| 716 | Acyl Desferrioxamin C15 (2) | 757.5753 | [M+H]+ | 757.5803 | 7 | - | **CCMSLIB00005716849** | 10.1128/mBio.00459-13 |
| 907 | Acyl Desferrioxamin C16 (2) | 771.5923 | [M+H]+ | 771.5959 | 5 | - | **CCMSLIB00005716850** | 10.1128/mBio.00459-13 |
| 1188 | Siomycin A (2) | 1648.4553 | [M+H]+ | 1648.4683 | 8 | - | **CCMSLIB00005716851** | 10.7164/antibiotics.22.364 |
| 11 | Siomycin A (2) | 824.7354 | [M+2H]2+ | 1648.4683 | 3 | - | **CCMSLIB00005716853** | 10.7164/antibiotics.22.364 |
| 18 | Siomycin B (2) | 1510.4183 | [M+H]+ | 1510.4254 | 5 | - | **CCMSLIB00005716852** | 10.7164/antibiotics.22.364 |
| 10 | Siomycin B (2) | 755.7147 | [M+2H]2+ | 1510.4254 | 2 | - | **CCMSLIB00005716854** | 10.7164/antibiotics.22.364 |
| 1000 | Siomycin C (2) | 832.2368 | [M+2H]2+ | 1663.4679 | 1 | - | **CCMSLIB00005716855** | 10.7164/antibiotics.22.364 |
| 339 | Siomycin D1 (2) | 817.7291 | [M+2H]2+ | 1634.4526 | 1 | - | **CCMSLIB00005716856** | 10.7164/antibiotics.33.1563 |
| 546 | Sphaericin (2) | 2157.116 | [M+1+H]+ | 2156.1116 | 1 | - | **CCMSLIB00005716857** | 10.1002/ejoc.201601334 |

[a]Feature ID coming from GNPS molecular networking job: *ID=92036537c21b44c29e509291e53f6382*

[b]Compound names are followed by classification (1 or 2) following CAWG guidelines.

[c]Difference between observed parent ion and reference ion (GNPS library or literature data) in ppm, abs.

[d]GNPS-generated similarity score, if applicable.

[e]Reference ID to entry in GNPS library. IDs in boldface have been deposited in the scope of this study.

[f]References to newly deposited entries. References starting with "10." indicate digital object identifiers (DOIs).

[g](4R)-4-((3R,5S,7R,9S,10S,13R,14S,15R,17R)-3,7,15-trihydroxy-10,13-dimethylhexadecahydro-1H-cyclopenta[a]phenanthren-17-yl)pentanoic acid.

Figure 4: Features identified in this study.

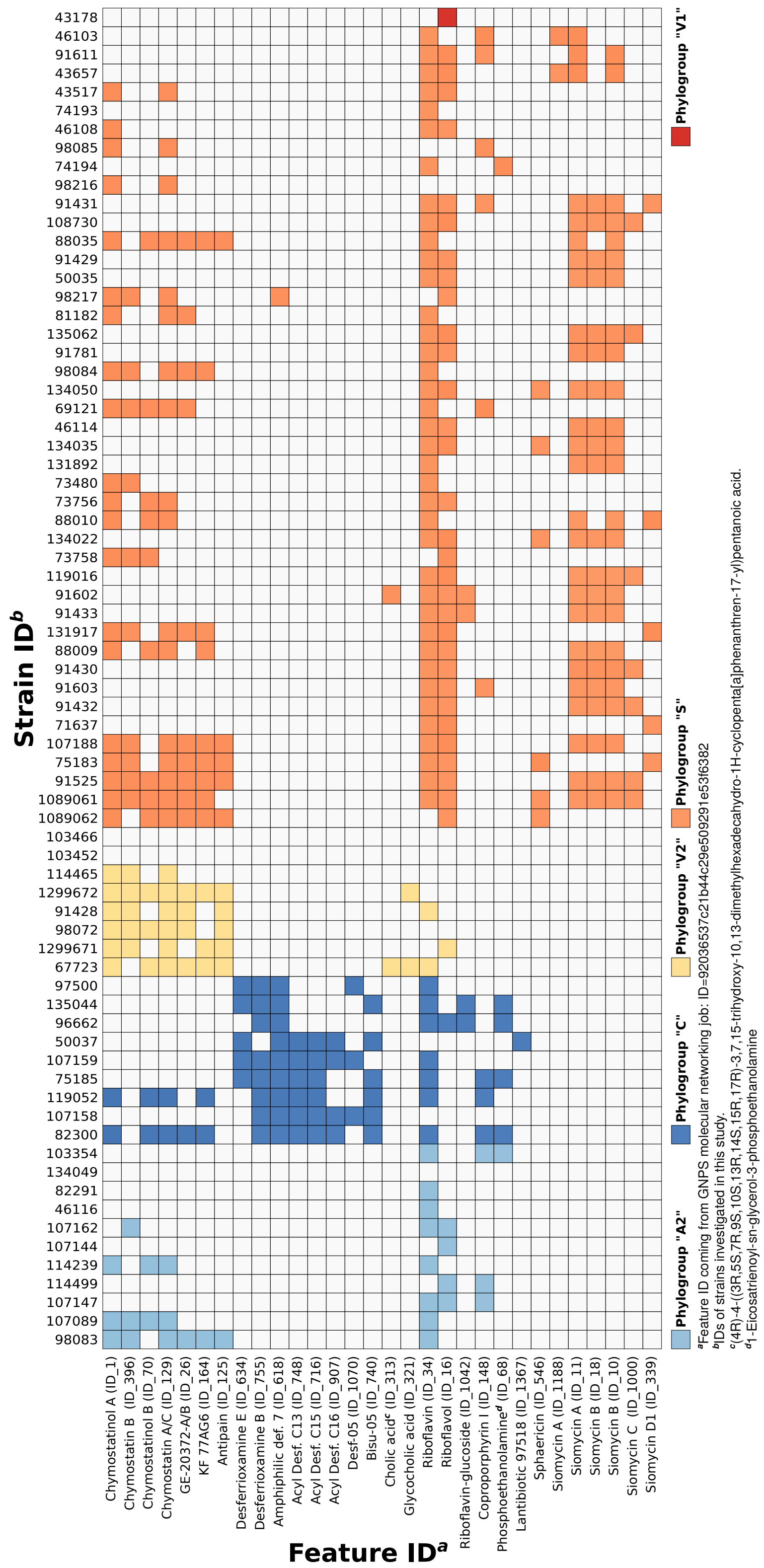

Figure 5: Table indicating presence/absence of identified features. Colours indicate the corresponding phylogroups, used to classify the strains cultivated in this study.

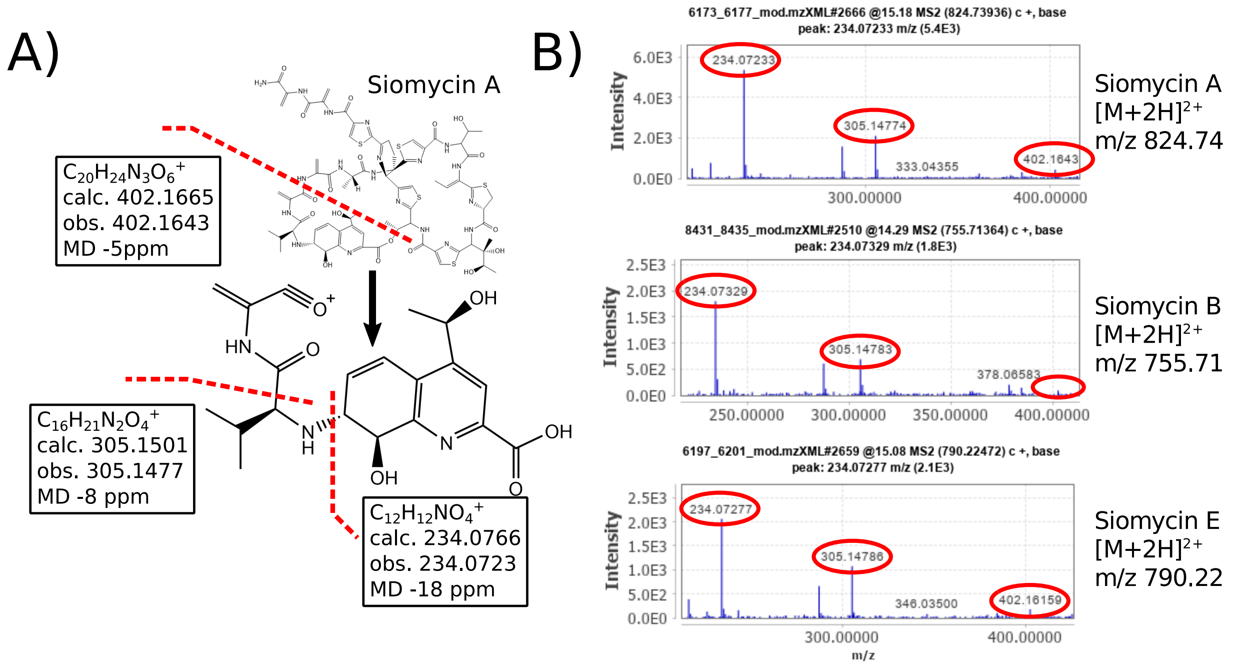

Figure 6: The quinalidic acid moiety of siomycin leads to characteristic fragmentation patterns. (A) shows a break between the $Ala^2$- and $Dha^3$ as well as $Thr^{12}$ and QA that leads to diagnostic peaks in $[M+2H]^{2+}$-ions. (B) shown are example spectra, observed in the present study.

A)

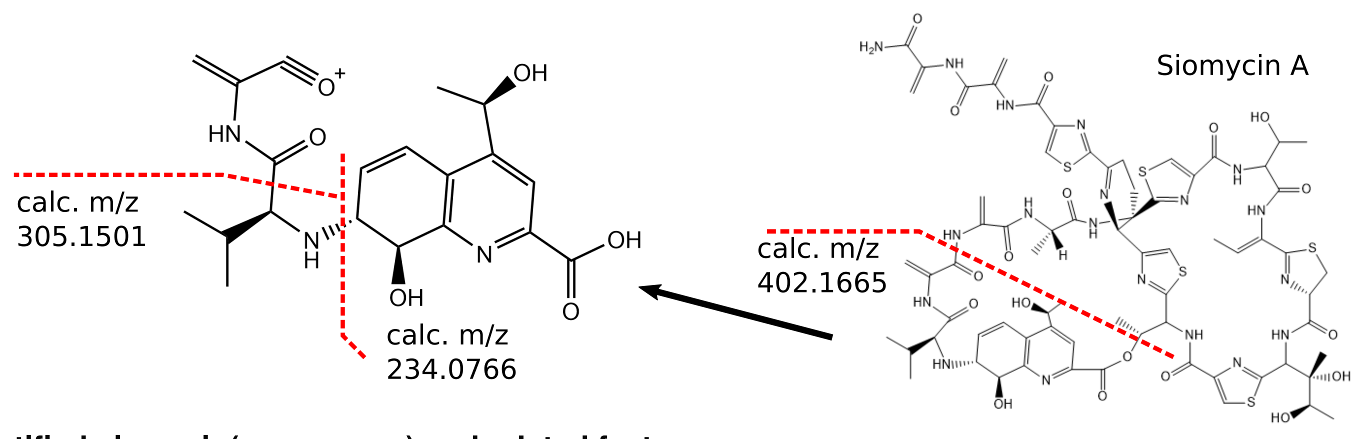

B)

**Identified siomycin(-congeners) and related features**

| Feature ID[a] | Name | Parent ion (*m/z*) | Par. ion adduct | Calc. mol. formula | Calc. par. ion (*m/z*) | *m/z* error[b] | Frag. *m/z* 234.08 | Frag. *m/z* 305.15 | Frag. *m/z* 402.17 | Comment |
|---|---|---|---|---|---|---|---|---|---|---|
| 11 | Siomycin A | 824.7354 | $[M+2H]^{2+}$ | $C_{71}H_{81}N_{19}O_{18}S_5$ | 1648.4683 | 3 | Yes | Yes | Yes | |
| 10 | Siomycin B | 755.7147 | $[M+2H]^{2+}$ | $C_{65}H_{75}N_{17}O_{16}S_5$ | 1510.4254 | 2 | Yes | Yes | Yes | |
| 1000 | Siomycin C | 832.2368 | $[M+2H]^{2+}$ | $C_{72}H_{82}N_{18}O_{19}S_5$ | 1663.4680 | 1 | Yes | Yes | Yes | |
| 339 | Siomycin D1 | 817.7291 | $[M+2H]^{2+}$ | $C_{70}H_{79}N_{19}O_{18}S_5$ | 1634.4527 | 1 | Yes* | Yes* | Yes* | *CH2 less on QA moiety |
| 9 | Siomycin E | 790.2255 | $[M+2H]^{2+}$ | $C_{68}H_{78}N_{18}O_{17}S_5$ | 1579.4474 | 2 | Yes | Yes | Yes | |
| 50 | Sio-like 2 | 906.2501 | $[M+2H]^{2+}$ | $C_{76}H_{90}N_{20}O_{21}S_6$ | 1811.4992 | 4 | Yes | Yes | Yes | Siom. A + N-Acetylcysteinly |
| 984 | Sio-like 3 | 885.2433 | $[M+2H]^{2+}$ | $C_{74}H_{88}N_{20}O_{20}S_6$ | 1769.4886 | 5 | Yes | Yes | Yes | Siom. A + N-Acetylcysteinly |
| 139 | Sio-like 4 | 914.2477 | $[M+2H]^{2+}$ | $C_{76}H_{90}N_{20}O_{22}S_6$ | 1827.4941 | 3 | Yes | Yes | Yes | Sio-like 2 + hydoxylation |
| 1168 | Sio-like 5 | 999.7384 | $[M+2H]^{2+}$ | - | *1998.4696* | - | Yes | Yes | Yes | *Multiple MFs possible* |
| 1466 | Sio-like 6 | 991.7438 | $[M+2H]^{2+}$ | - | *1982.4804* | - | Yes | Yes | Yes | *Multiple MFs possible* |
| 1064 | Sio-like 7 | 878.7283 | $[M+2H]^{2+}$ | - | *1756.4494* | - | Yes | Yes | Yes | *Multiple MF possible* |
| 1358 | Sio-like 8 | 987.7658 | $[M+2H]^{2+}$ | $C_{81}H_{99}N_{21}O_{24}S_7$ | 1974.5295 | 3 | Yes | Yes | Yes | Siom. A + 2xN-Acetylcysteinly |

[a]Feature ID from GNPS mol. networking job: *ID=92036537c21b44c29e509291e53f6382*
[b]Difference between observed parent ion and $[M+H]^+$ of calculated molecular formula in ppm, abs.

Figure 7: List of identified siomycins, congeners and related features, showing similar tandem mass fragmentation pattern.

Siomycin E
$[M+H]^+$
$C_{68}H_{79}N_{18}O_{17}S_5^+$
calc: 1579.4474
obs: 1579.4428
MD: 3ppm

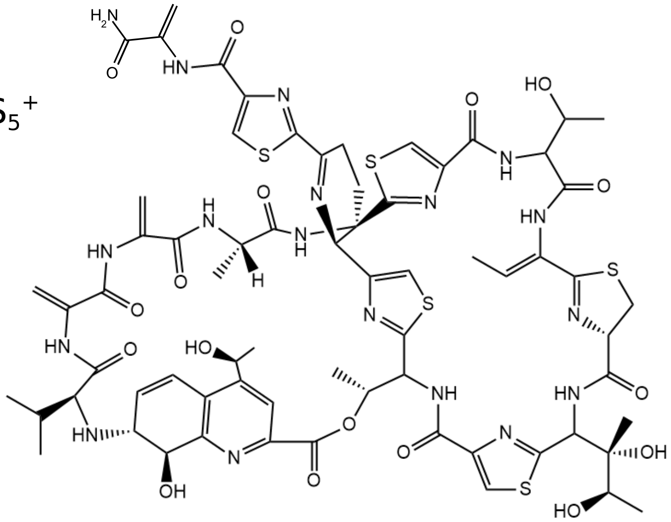

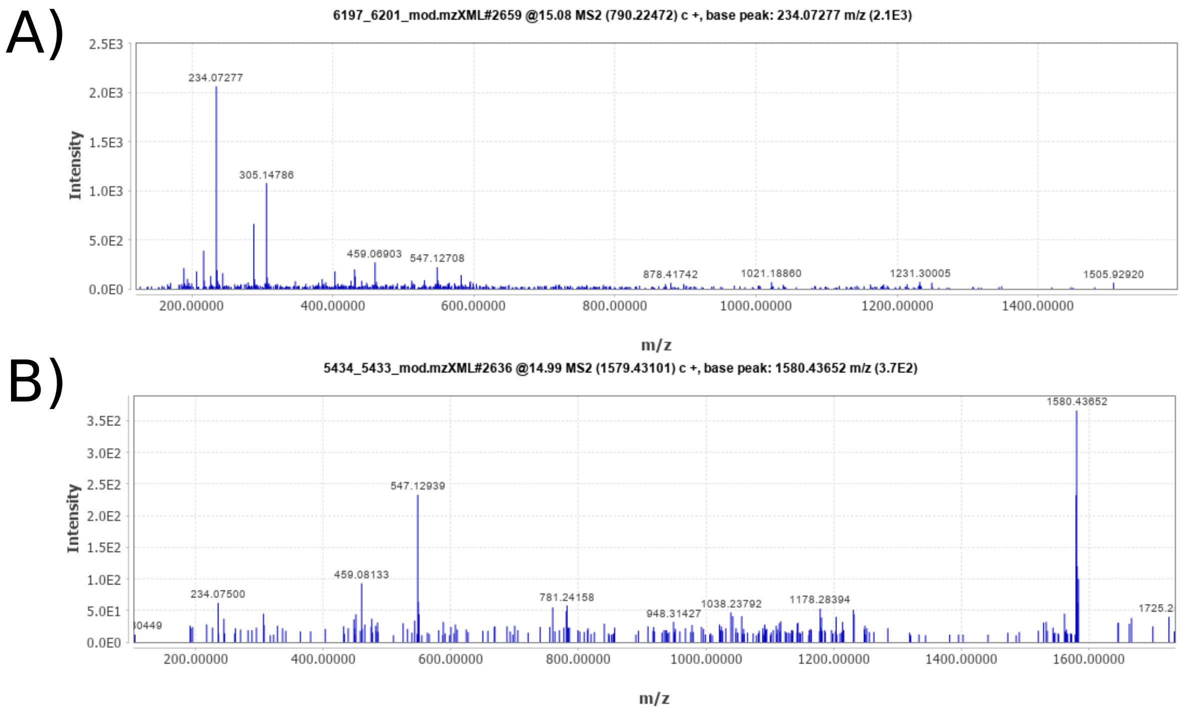

Figure 8: MS$^2$-fragmentation spectra of the $[M+H]^+$ (A) and $[M+2H]^{2+}$ (B) ions of siomycin E.

| | Leader Peptide | Core Peptide | |
|---|---|---|---|
| **TsrH** | MSNAAL - - - E I GVEGLTGLDVDTLE I SDYMDETLLDGEDLTVTM | I ASASCTTC I CTCSCSS | Liao et al. Cell Chemical Biology 2009 |
| **SioH** | MSTAA I VGQE I GVDGLTGLDVDALE I SDYMDETLLDGEDLSVTM | VSSASCTTC I CTCSCSS | Liao et al. Cell Chemical Biology 2009 |
| **RiPP4 (ID91781)** | MSTAT - - SQS I GVESLTGLDVDMLE I SDY I DETLLDTADLTVTM | VSSASCTTC I CTCSCSS | *This study* |
| **TpnA** | MSAPT - E I QNLGVVGLTGLDTDTLE I SDYLDESLLDEHDLTVTM | VASASCTTC I CTCSCSS | Ichikawa et al. J. Am. Chem. Soc. 2018 |

Figure 9: Comparison of thiopeptide precursor peptides. TsrH = thiostrepton precursor peptide; SioH = siomycin precursor peptide; TpnA = thiopeptin precursor peptide; RiPP4 = precursor peptide of siomycin in the genome of strain ID91781, detected in this study. All thiopeptide core peptides have two serines at their C-terminal end, which are eventually modified to dehydroalanines.

#### Annotated MS2LDA-motif list

| Name | Degree[a] | Annotation |
|---|---|---|
| motif_191 | 153 | Desferrioxamine-related motif |
| motif_104 | 104 | Desferrioxamine-related motif |
| motif_474 | 66 | Siomycin-related motif |
| motif_392 | 54 | Putative siderophore-related motif |
| motif_449 | 50 | Desferrioxamine-related motif |
| motif_47 | 49 | Chymostatin-reated motif |
| motif_385 | 46 | Desferrioxamine-related motif |
| motif_134 | 46 | Chymostatin-reated motif |
| motif_217 | 45 | Desferrioxamine-related motif |
| motif_363 | 45 | Siomycin-related motif |
| motif_93 | 40 | Siomycin-related motif |
| motif_111 | 37 | Desferrioxamine-related motif |
| motif_506 | 37 | Desferrioxamine-related motif |
| motif_147 | 35 | Chymostatin-related motif |
| motif_533 | 32 | Chymostatin-related motif |
| motif_174 | 29 | Desferrioxamine-related motif |
| motif_530 | 27 | Chymostatin-related motif |
| motif_408 | 22 | Chymostatin-related motif |
| motif_154 | 22 | Siomycin-related motif |
| motif_412 | 22 | Desferrioxamine-related motif |
| motif_529 | 20 | Desferrioxamine-related motif |
| motif_102 | 20 | Siomycin-related motif |
| motif_434 | 19 | Putative siderophore-related motif |
| motif_259 | 18 | Desferrioxamine-related motif |
| motif_203 | 18 | Putative siderophore-related motif |
| motif_285 | 17 | Siomycin-related motif |
| motif_364 | 15 | Siomycin-related motif |
| motif_192 | 15 | Sphaericin-related motif |
| motif_256 | 14 | Desferrioxamine-related |
| motif_301 | 13 | Siomycin-related motif |

[a]Number of features, associated to a particular motifs.
Note: the same feature can be present in multiple motifs.

Figure 10: Annotated MS2LDA-motifs, identified in this study.

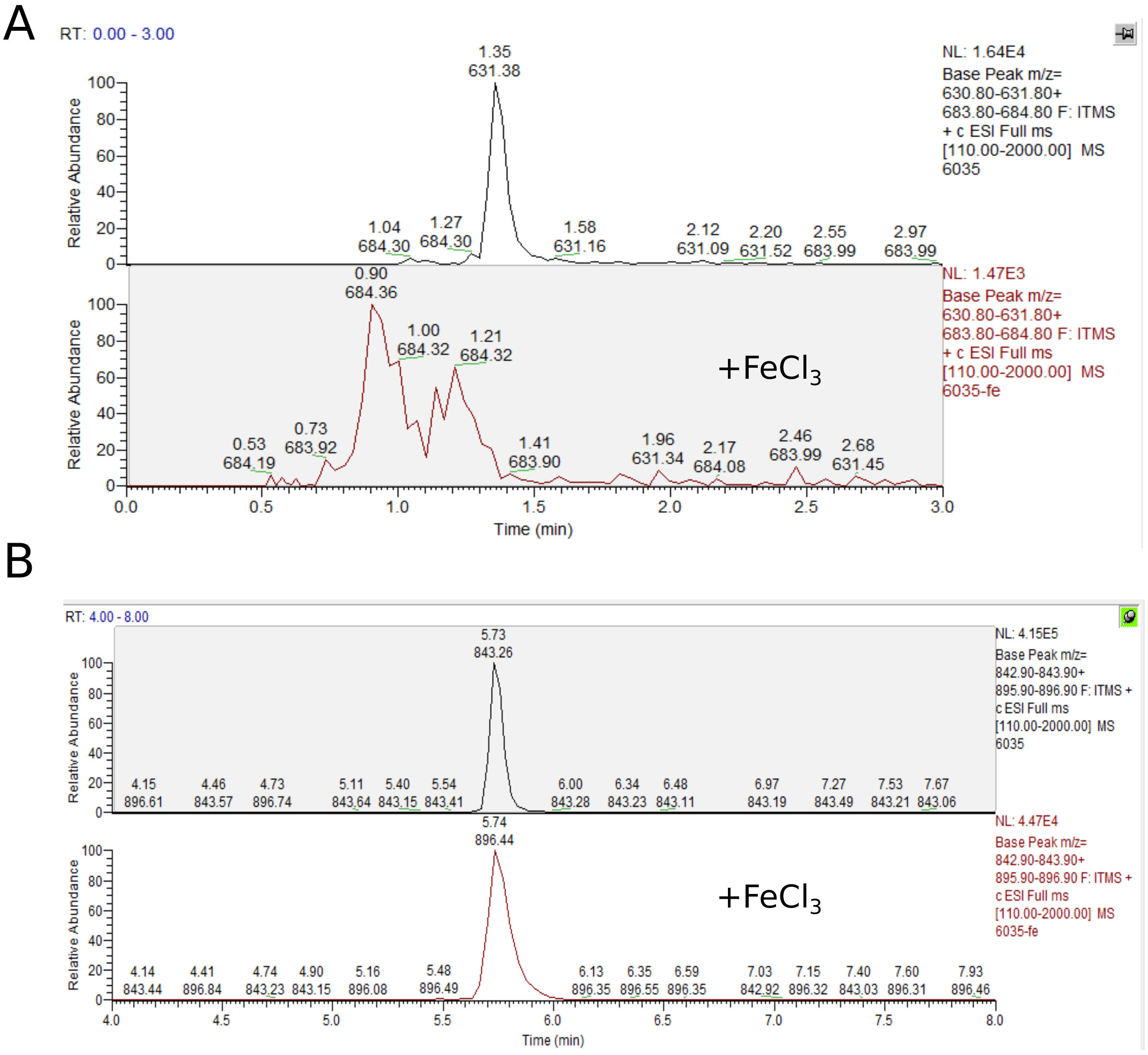

Figure 11: Iron-complexion of metabolite $m/z$ 631.4 and its related metabolite $m/z$ 843.47 upon addition of FeCl3 (original LCMS-traces of examples shown in Figure 10 of the main text).

#### Annotation of planochelin-related features

| Feature ID[a] | Molecular family | Parent ion (*m/z*) | Par. ion adduct | Calc. mol. formula | Calc. par. ion (*m/z*) | *m/z* error (ppm)[b] | Phylo-group | Producing strains |
|---|---|---|---|---|---|---|---|---|
| 279 | I | 631.3408 | $[M+H]^+$ | $C_{26}H_{46}N_8O_{10}$ | 631.3415 | 1 | V2 | 67723,1299671,1299672,98072,91428 |
| 282 | I | 757.4677 | $[M+H]^+$ | $C_{35}H_{64}N_8O_{10}$ | 757.4824 | 20 | V2 | 67723,1299671,98072 |
| 268 | I | 715.4325 | $[M+H]^+$ | $C_{32}H_{58}N_8O_{10}$ | 715.4354 | 4 | V2 | 67723,1299671,98072 |
| 1547 | I | 771.4933 | $[M+H]^+$ | $C_{36}H_{66}N_8O_{10}$ | 771.4980 | 5 | V2 | 1299671 |
| 262 | II | 829.4587 | $[M+H]^+$ | $C_{37}H_{64}N_8O_{13}$ | 829.4671 | 10 | V2, S | 67723,1299671,1089062,1089061 |
| 267 | II | 843.4772 | $[M+H]^+$ | $C_{38}H_{66}N_8O_{13}$ | 843.4827 | 6 | V2, S | 67723,1299671,1299672,1089062,98072,1089061 |
| 275 | II | 815.4517 | $[M+H]^+$ | $C_{36}H_{62}N_8O_{13}$ | 815.4515 | 1 | V2 | 67723 |
| 330 | II | 871.5077 | $[M+H]^+$ | $C_{40}H_{70}N_8O_{13}$ | 871.5141 | 7 | V2 | 67723,1299671 |
| 239 | II | 841.4616 | $[M+H]^+$ | $C_{38}H_{64}N_8O_{13}$ | 841.4671 | 7 | V2, S | 67723,1089061 |
| 324 | II | 813.4680 | $[M+H]^+$ | $C_{36}H_{60}N_8O_{13}$ | 813.3458 | 40 | V2 | 67723 |
| 326 | II | 869.4919 | $[M+H]^+$ | $C_{40}H_{68}N_8O_{13}$ | 869.4984 | 8 | V2 | 67723 |
| 273 | II | 857.4919 | $[M+H]^+$ | $C_{39}H_{68}N_8O_{13}$ | 857.4984 | 8 | V2, S | 1089061,67723,1299671,98072 |
| 1552 | III | 728.4164 | $[M+H]^+$ | $C_{35}H_{53}N_9O_8$ | 728.4095 | 9 | V2 | 1299671 |
| 1549 | III | 742.4308 | $[M+H]^+$ | $C_{36}H_{55}N_9O_8$ | 742.4252 | 8 | V2 | 1299671 |
| 1553 | III | 714.4003 | $[M+H]^+$ | $C_{34}H_{51}N_9O_8$ | 714.3939 | 9 | V2 | 1299671 |
| 1556 | III | 756.4452 | $[M+H]^+$ | $C_{37}H_{58}N_9O_8$ | 756.4408 | 6 | V2 | 1299671 |
| 314 | IV | 1067.5659 | $[M+H]^+$ | $C_{48}H_{78}N_{10}O_{17}$ | 1067.5625 | 3 | V2 | 67723 |
| 289 | IV | 905.5136 | $[M+H]^+$ | $C_{42}H_{68}N_{10}O_{12}$ | 905.5096 | 4 | V2 | 67723,1299671,98072 |
| 1222 | IV | 919.5277 | $[M+H]^+$ | $C_{43}H_{70}N_{10}O_{12}$ | 919.5253 | 2 | V2 | 1299671,98072 |
| 298 | IV | 891.4984 | $[M+H]^+$ | $C_{41}H_{66}N_{10}O_{12}$ | 891.4940 | 5 | V2 | 67723,1299671,98072 |
| 288 | IV | 877.4821 | $[M+H]^+$ | $C_{40}H_{64}N_{10}O_{12}$ | 877.4789 | 4 | V2 | 67723,1299671,98072 |

[a]Feature ID from GNPS mol. networking job: *ID=92036537c21b44c29e509291e53f6382*
[b]Difference between observed parent ion and $[M+H]^+$ of calc. mol. formula in ppm, abs.

Figure 12: Overview of putative siderophores identified in this study.

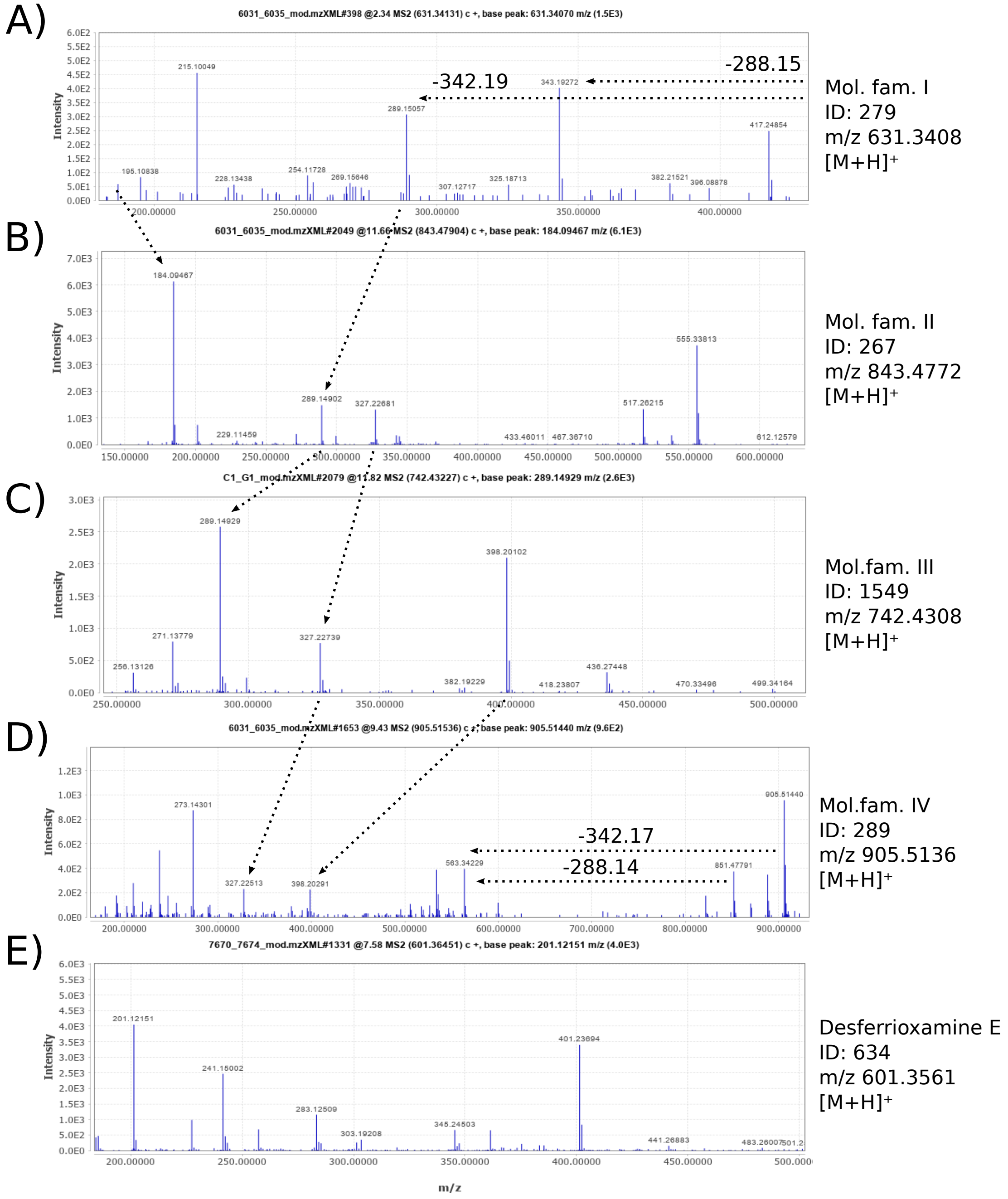

Figure 13: Examples for similarities in tandem mass fragmentation between iron-complexing metabolites from different molecular families, both in common fragments and neutral losses. (A) feature ID279 (*m/z* 631.3), representative of molecular family I. (B) feature ID267 (*m/z* 843.5), representative of molecular family II. (C) feature ID1549 (*m/z* 742.4), representative of molecular family III. (D) feature ID289 (*m/z* 905.5), representative of molecular family IV. (E) desferrioxamine E, which shows no similarities to features A-D.

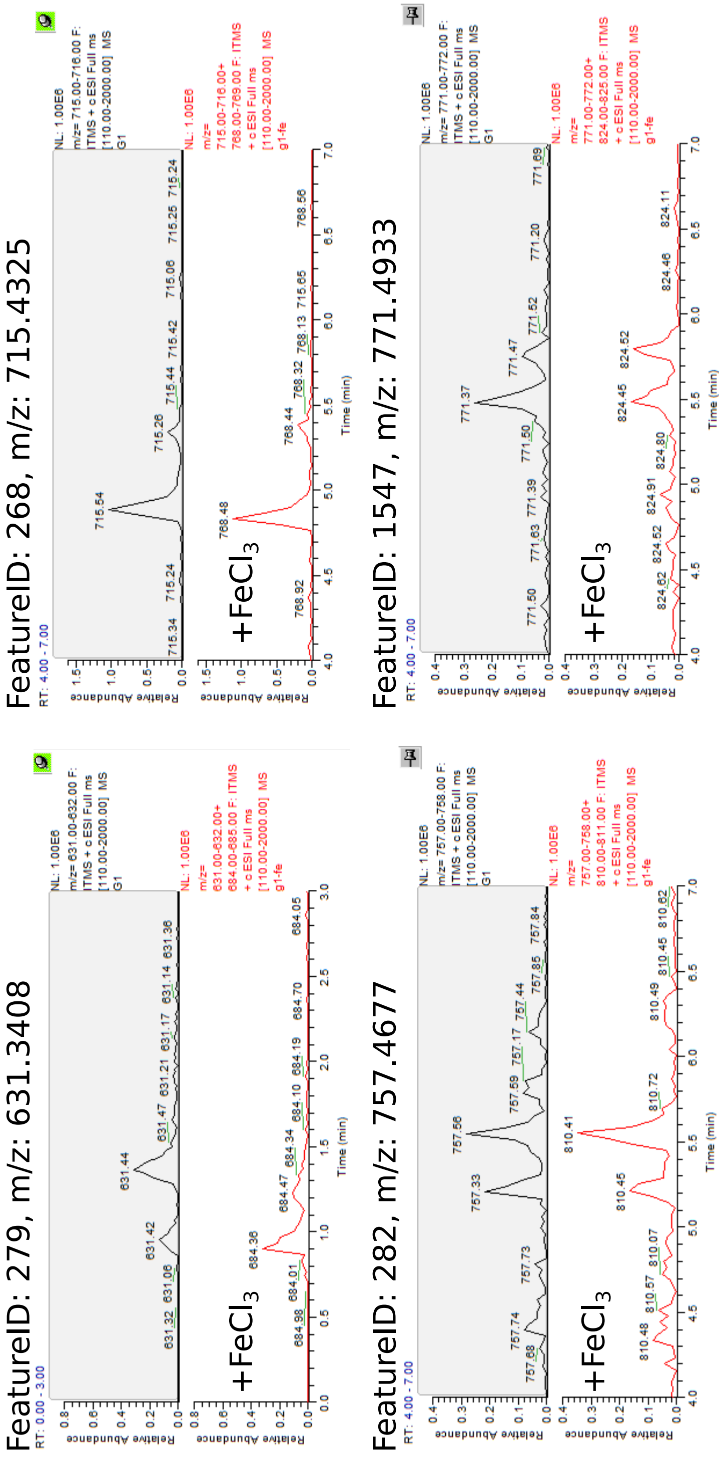

Figure 14: Iron-complexion of feature ID279 (631.3 *m/z*) and its related metabolites in molecular family I.

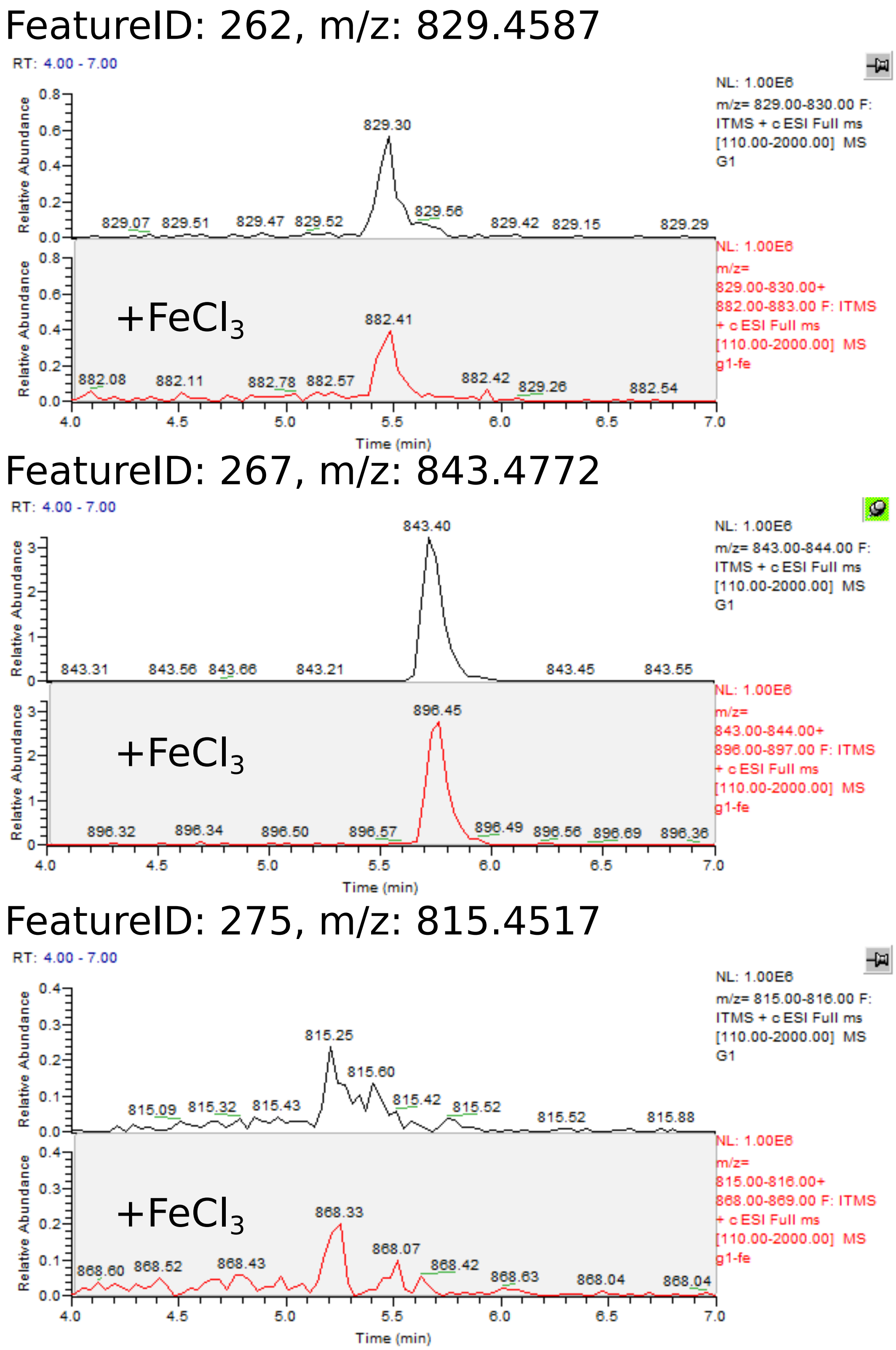

Figure 15: Iron-complexion of putative siderophore metabolites in molecular family II.

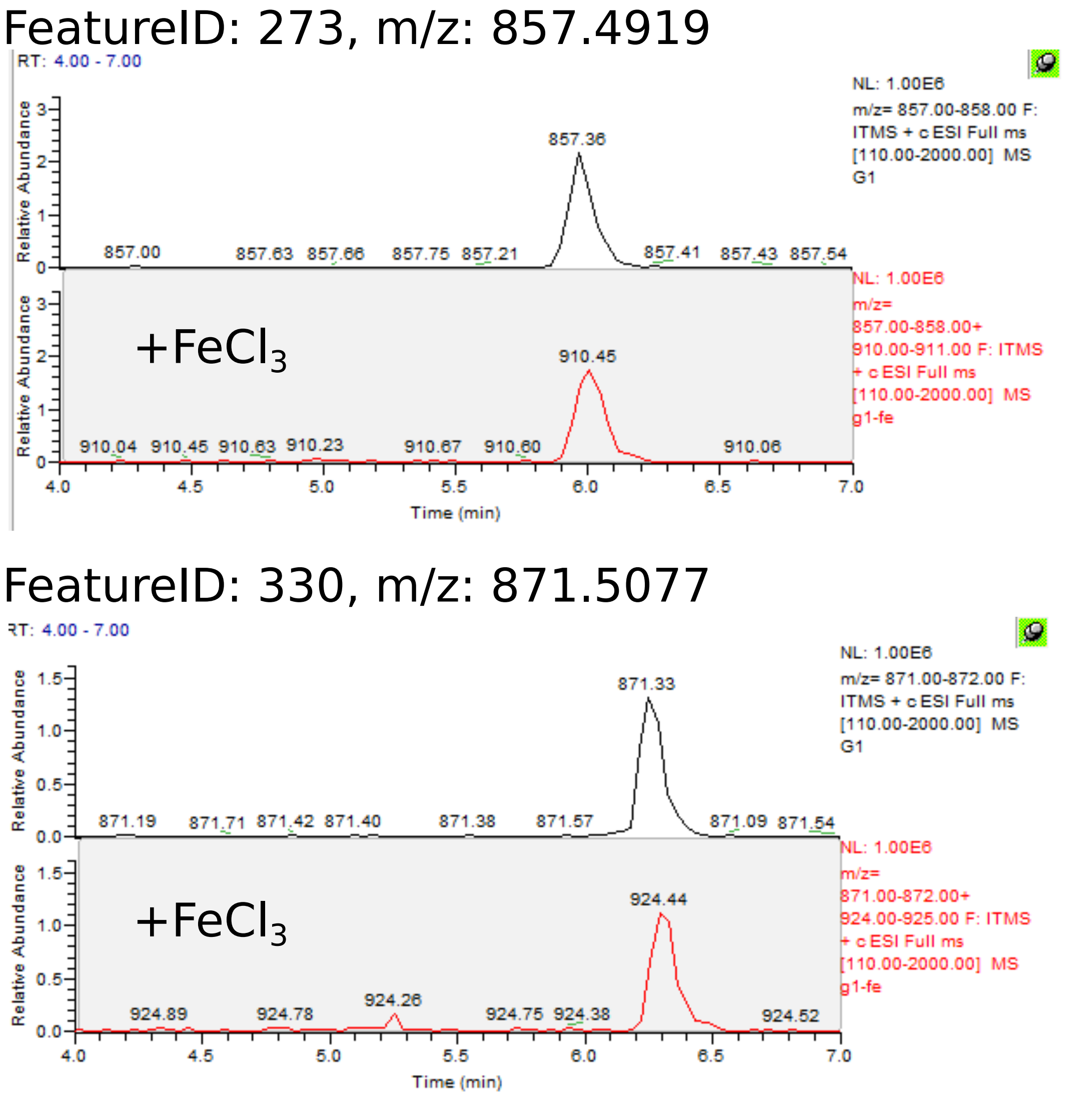

Figure 16: Iron-complexion of putative siderophore metabolites in molecular family II.

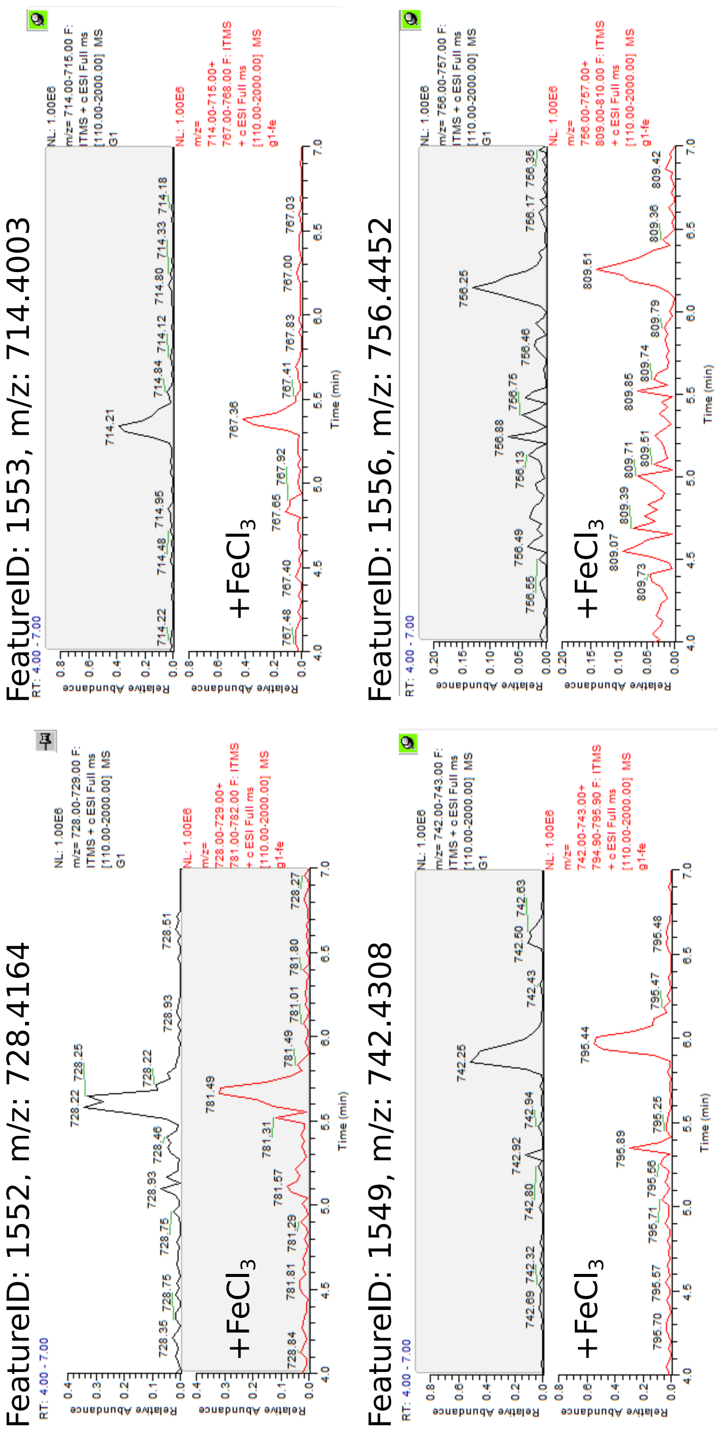

Figure 17: Iron-complexion of putative siderophore metabolites in molecular family III.

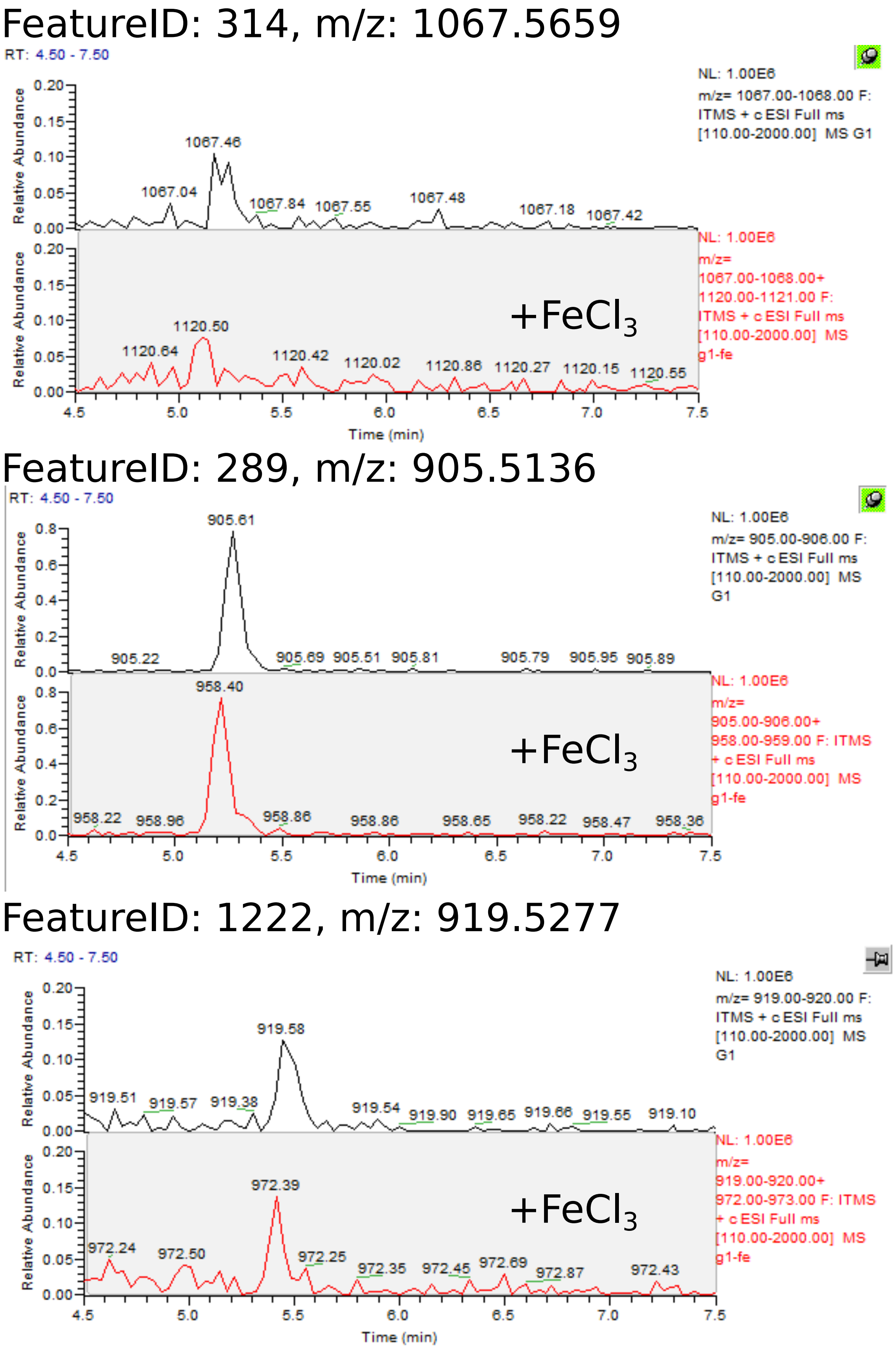

Figure 18: Iron-complexion of putative siderophore metabolites in molecular family IV.

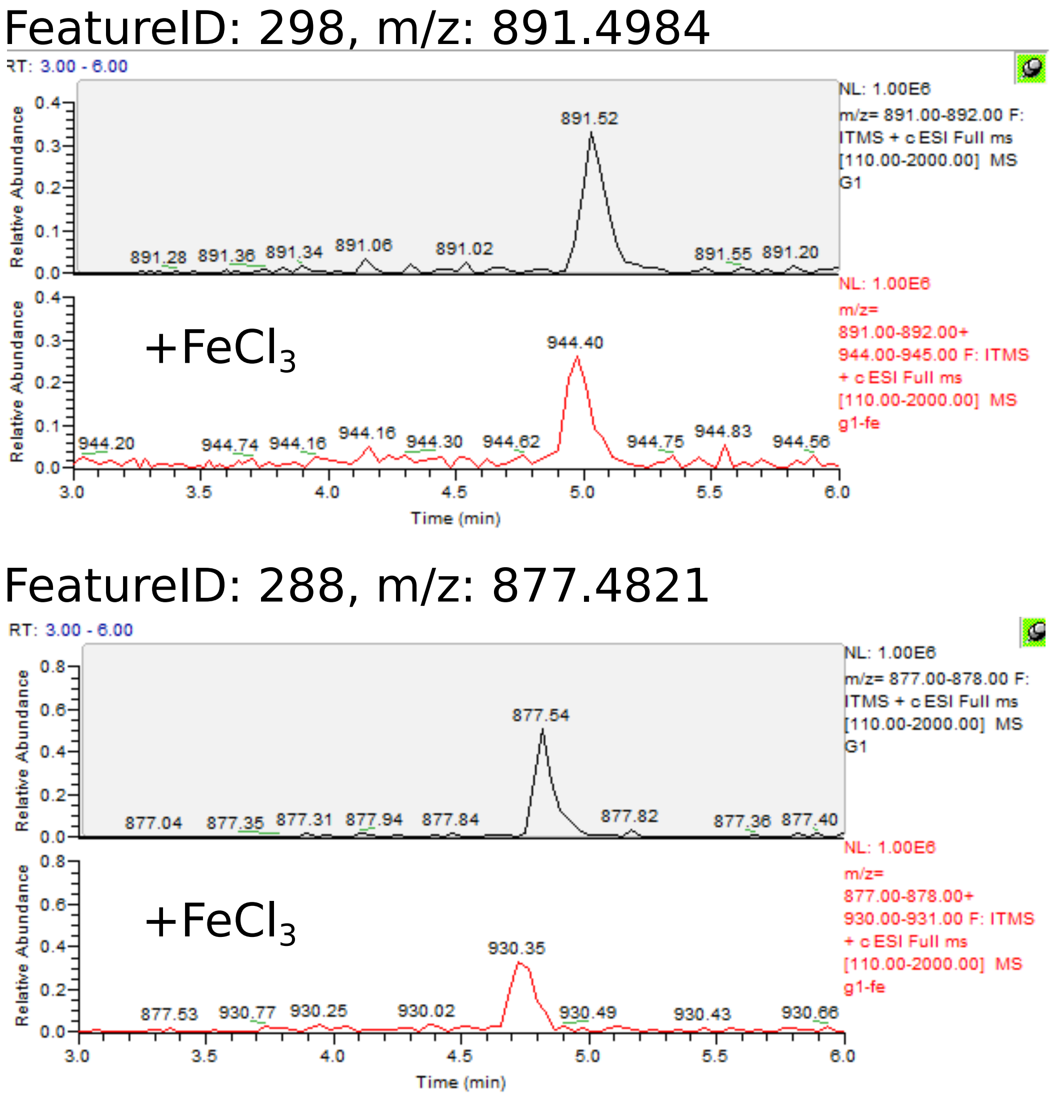

Figure 19: Iron-complexion of putative siderophore metabolites in molecular family IV.

Antipain
Found: 303.18
Calc: 303.18
Δppm: 3

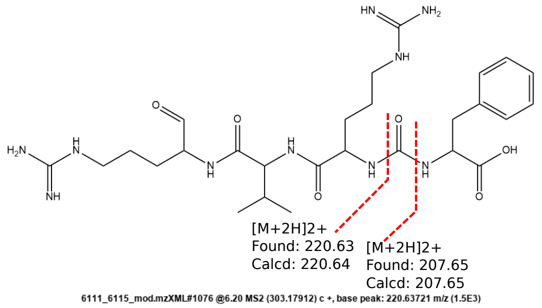

A)

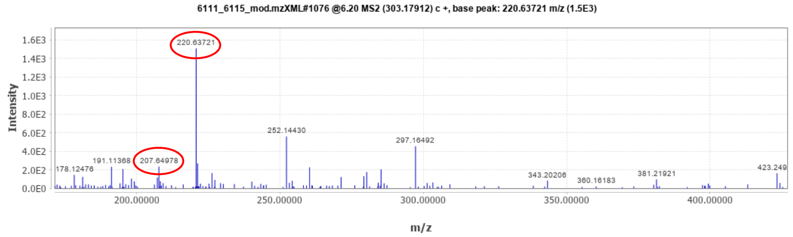

ESI-MS/MS spectrum for the [M+2H]2+ ion of **antipain**, measured on a Bruker micrOTOF-Q III instrument, detected in the extracts 6111 and **6115**. Peaks circled in red are consistent with the fragmentation above.

B)

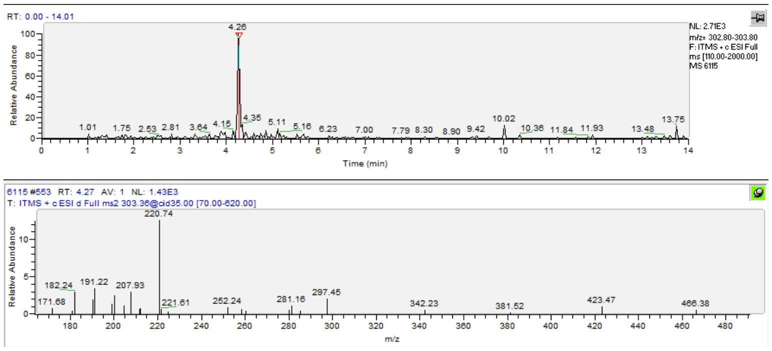

ESI-MS/MS spectrum for the [M+2H]2+ ion of **antipain** measured on a ThermoFisher LTQ linear ion trap instrument, detected in the extract **6115**. The main MS2 fragments correspond to the ones detected on the high resolution instrument (A). The retention time and the main MS2 fragments correspond to the ones detected for the antipain standard (C).

C)

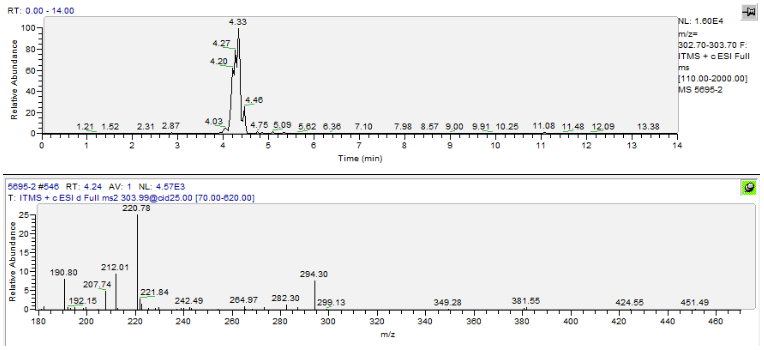

ESI-MS/MS spectrum for the [M+2H]2+ ion of **antipain**, measured on a ThermoFisher LTQ linear ion trap instrument, from an authentic, in-house characterized standard of **antipain**.

Figure 20: Proof for the annotation of the metabolite antipain.

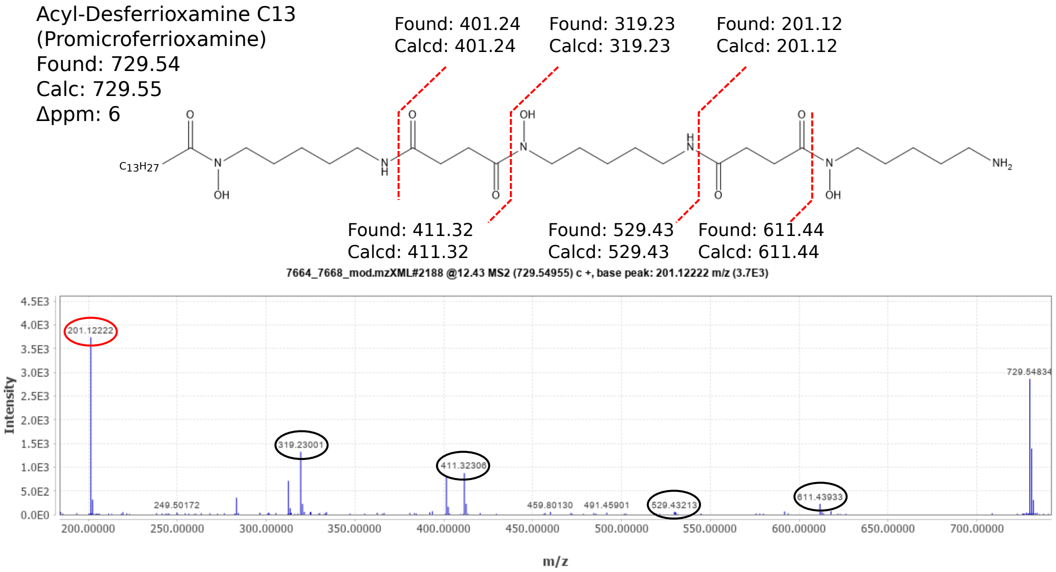

ESI-MS/MS spectrum for the [M+H]+ ion of **Acyl-Desferrioxamine C13 (Promicroferrioxamine)**, measured on a Bruker micrOTOF-Q III instrument. Peaks circled in black are consistent with the published spectrum (Yang et al. Angew Chem Int Ed 50.26(2011):5839-5842), peaks circled in red are consistent with the fragmentation pattern indicated above.

Figure 21: Proof for the annotation of metabolite acyl-desferrioxamine C13.

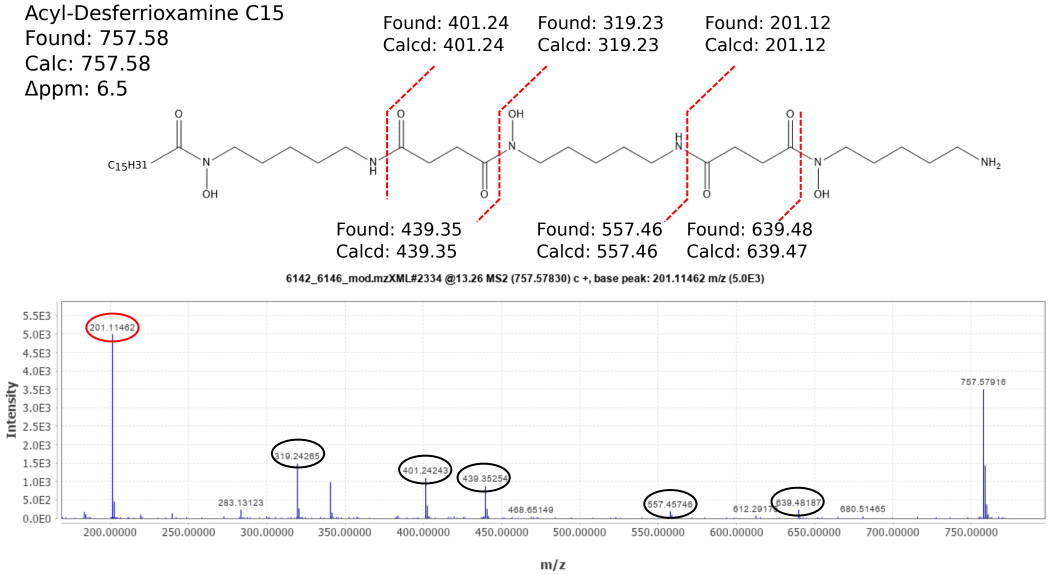

ESI-MS/MS spectrum for the [M+H]+ ion of **Acyl-Desferrioxamine C15,** measured on a Bruker micrOTOF-Q III instrument. Peaks circled in black are consistent with the published spectrum (Traxler et al. MBio 4.4(2013): e00459-13), peaks circled in red are consistent with the fragmentation pattern indicated above.

Figure 22: Proof for the annotation of metabolite acyl-desferrioxamine C15.

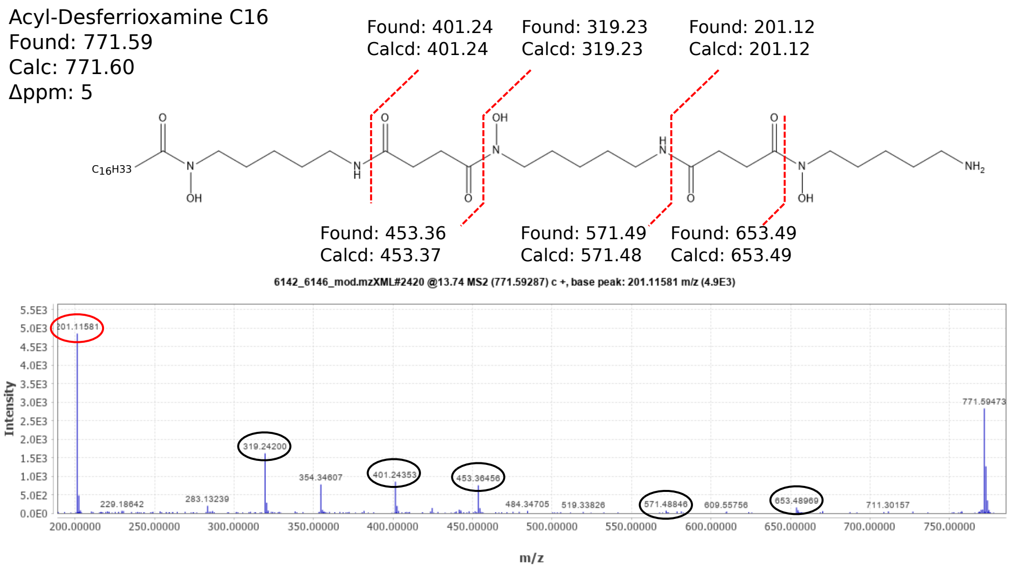

ESI-MS/MS spectrum for the [M+H]+ ion of **Acyl-Desferrioxamine C16,** measured on a Bruker micrOTOF-Q III instrument. Peaks circled in black are consistent with the published spectrum (Traxler et al. MBio 4.4(2013): e00459-13), peaks circled in red are consistent with the fragmentation pattern indicated above.

Figure 23: Proof for the annotation of metabolite acyl-desferrioxamine C16.

Chymostatin A/C
Found: 608.31
Calc: 608.32
Δppm: 9

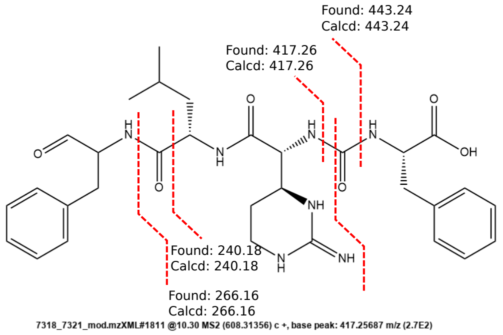

A)

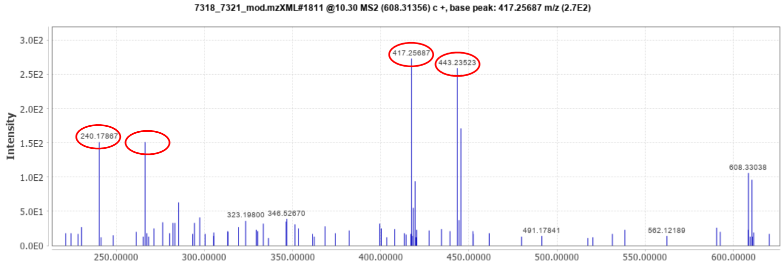

ESI-MS/MS spectrum for the [M+H]+ ion of **chymostatin A/C**, measured on a Bruker micrOTOF-Q III instrument, detected in the extracts **7318** and 7321. Peaks circled in red are consistent with the fragmentation above.

B)

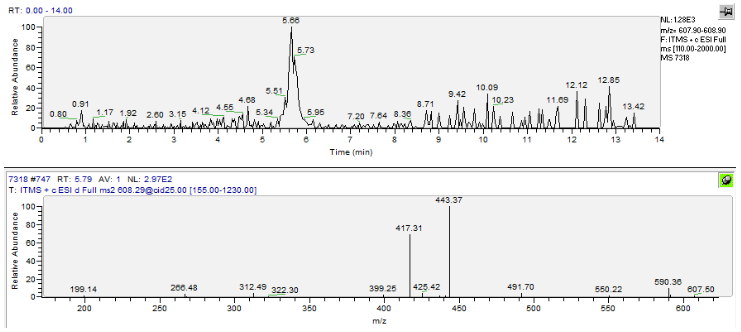

ESI-MS/MS spectrum for the [M+H]+ ion of **chymostatin A/C** measured on a ThermoFisher LTQ linear ion trap instrument, detected in the extract **7318**. The main MS2 fragments correspond to the ones detected on the high resolution instrument (A). The retention time and the main MS2 fragments correspond to the ones detected for the chymostatin standard (C).

C)

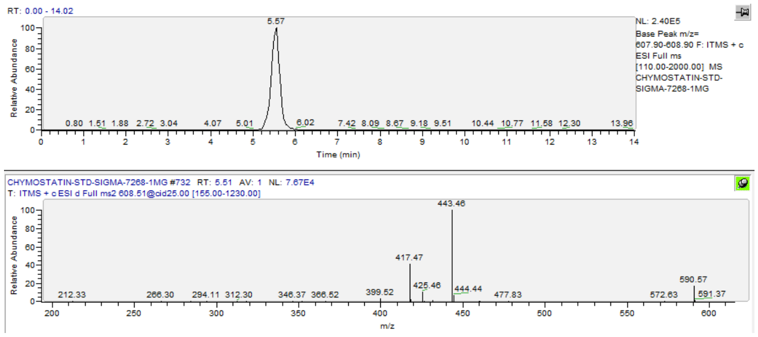

ESI-MS/MS spectrum for the [M+H]+ ion of **chymostatin A/C**, measured on a ThermoFisher LTQ linear ion trap instrument, from **Chymostatin Standard** of Sigma Aldrich (Prod. Nr. C7268, CAS 9076-44-2).

Figure 24: Proof for the annotation of metabolite chymostatin A or C (Leu/Ile)

A)

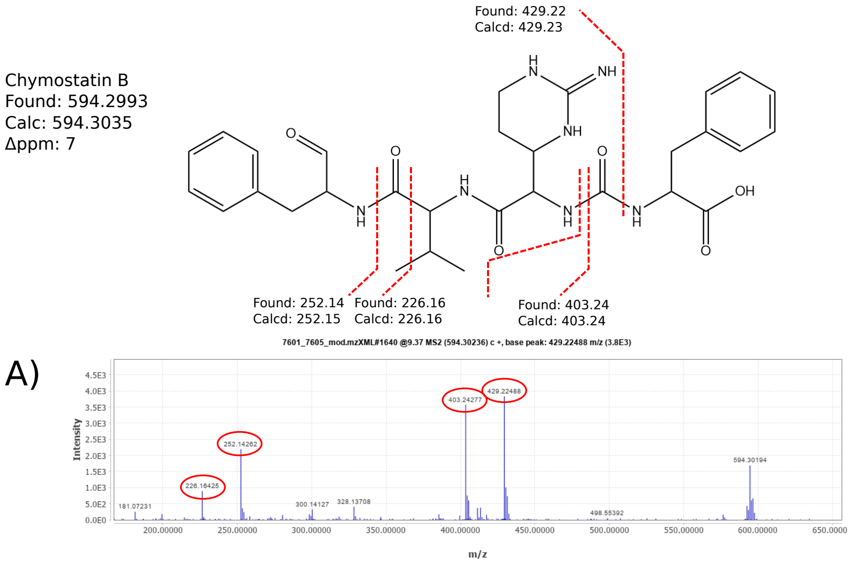

ESI-MS/MS spectrum for the [M+H]+ ion of **chymostatin B**, measured on a Bruker micrOTOF-Q III instrument, detected in the extracts 7601 and **7605**. Peaks circled in red are consistent with the fragmentation above.

B)

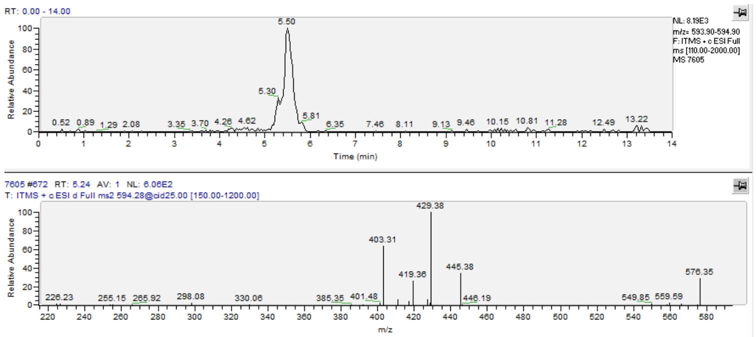

ESI-MS/MS spectrum for the [M+H]+ ion of **chymostatin B** measured on a ThermoFisher LTQ linear ion trap instrument, detected in the extract **7605**. The main MS2 fragments correspond to the ones detected on the high resolution instrument (A). The retention time and the main MS2 fragments correspond to the ones detected for the chymostatin standard (C).

C)

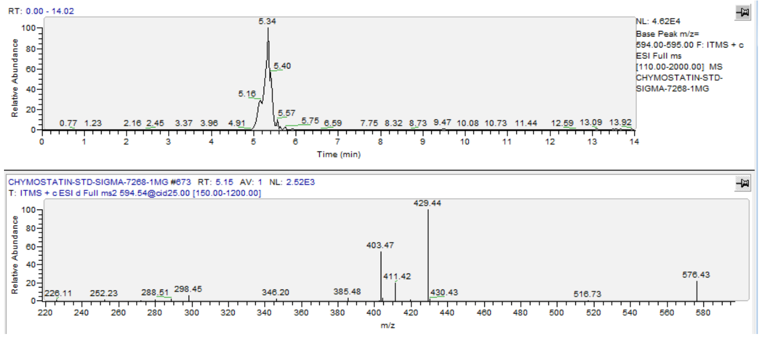

ESI-MS/MS spectrum for the [M+H]+ ion of **chymostatin B**, measured on a ThermoFisher LTQ linear ion trap instrument, from **Chymostatin Standard** of Sigma Aldrich (Prod. Nr. C7268, CAS 9076-44-2).

Figure 25: Proof for the annotation of metabolite chymostatin B.

Chymostatinol A
Found: 596.3151
Calc: 596.3191
Δppm: 6.5

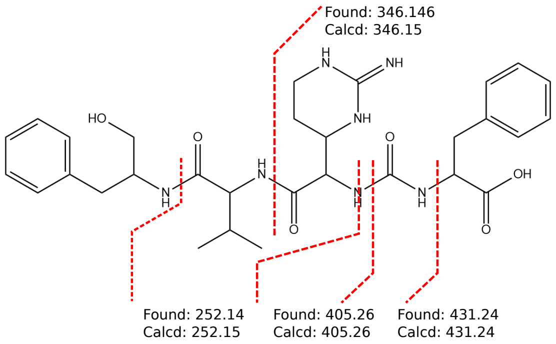

A)

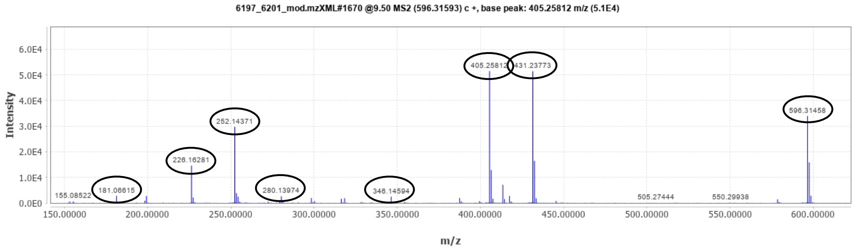

ESI-MS/MS spectrum for the [M+H]+ ion of **chymostatinol A**, measured on a Bruker micrOTOF-Q III instrument, detected in the extracts 6197 and 6201. Peaks circled in black are consistent with the published fragmentation spectrum (GNPS:CCMSLIB00000577828).

B)

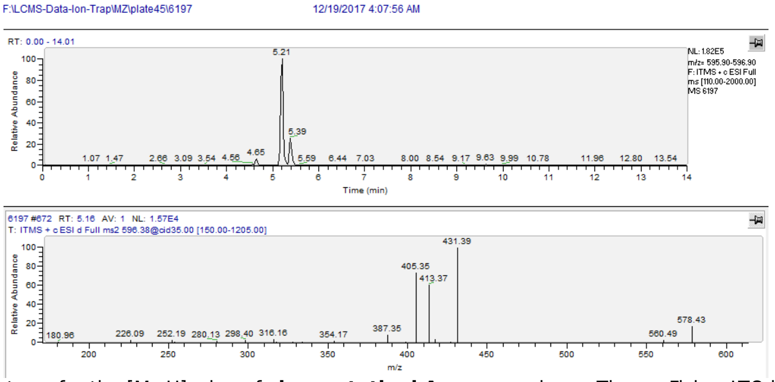

ESI-MS/MS spectrum for the [M+H]+ ion of **chymostatinol A**, measured on a ThermoFisher LTQ linear ion trap instrument, detected in the extract 6197. The main MS2 fragments correspond to the ones detected on the high resolution instrument (A). The retention time and the main MS2 fragments correspond to the ones detected for the chymostatin standard (C).

C)

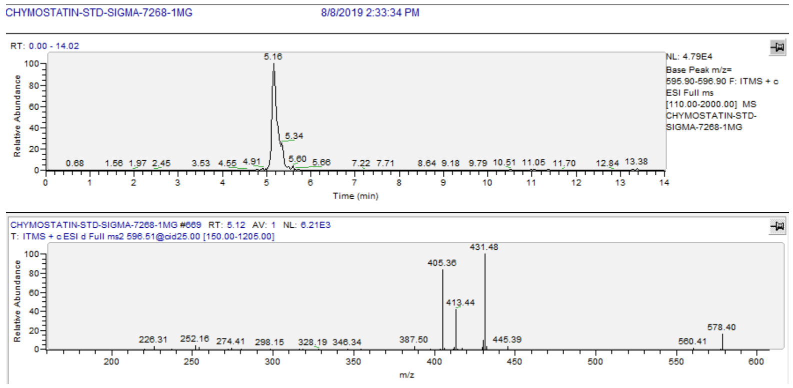

ESI-MS/MS spectrum for the [M+H]+ ion of **chymostatinol A**, measured on a ThermoFisher LTQ linear ion trap instrument, from **Chymostatin Standard** of Sigma Aldrich (Prod. Nr. C7268, CAS 9076-44-2).

Figure 26: Proof for the annotation of metabolite chymostatinol A.

Chymostatinol B
Found: 610.33
Calc: 610.35
Δppm: 8

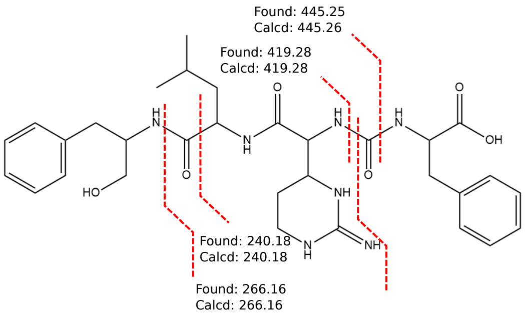

A)

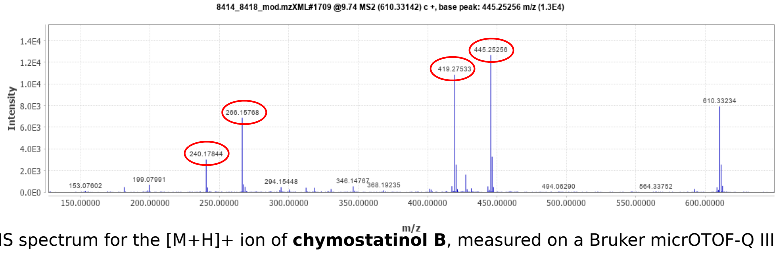

ESI-MS/MS spectrum for the [M+H]+ ion of **chymostatinol B**, measured on a Bruker micrOTOF-Q III instrument, detected in the extracts 8414 and **8418**. Peaks circled in red are consistent with the fragmentation above.

B)

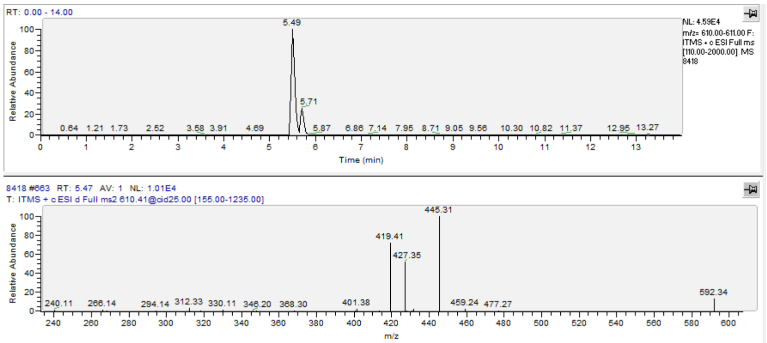

ESI-MS/MS spectrum for the [M+H]+ ion of **chymostatinol B** measured on a ThermoFisher LTQ linear ion trap instrument, detected in the extract **8418**. The main MS2 fragments correspond to the ones detected on the high resolution instrument (A). The retention time and the main MS2 fragments correspond to the ones detected for the chymostatin standard (C).

C)

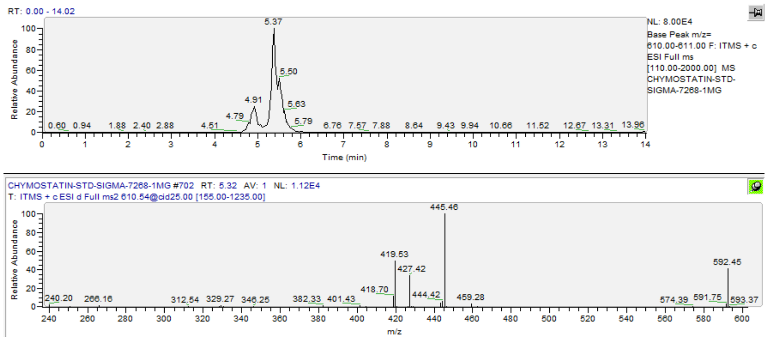

ESI-MS/MS spectrum for the [M+H]+ ion of **chymostatinol B**, measured on a ThermoFisher LTQ linear ion trap instrument, from **Chymostatin Standard** of Sigma Aldrich (Prod. Nr. C7268, CAS 9076-44-2).

Figure 27: Proof for the annotation of metabolite chymostatinol B.

GE-20372-A/B
Found: 612.31
Calc: 612.31
Δppm: 6

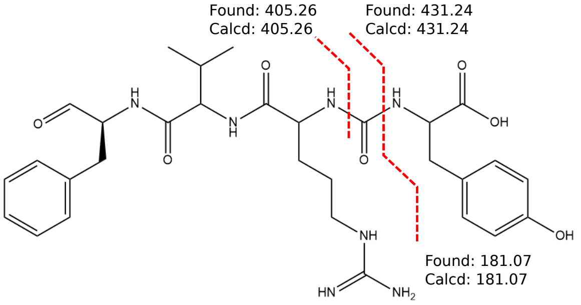

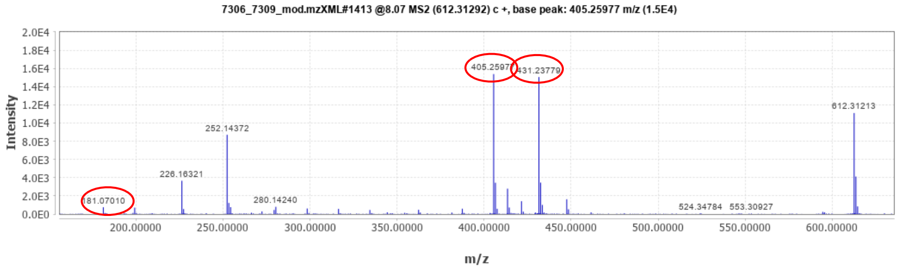

ESI-MS/MS spectrum for the [M+H]+ ion of **GE-20372-A/B**, measured on a Bruker micrOTOF-Q III instrument. Peaks circled in red are consistent with the fragmentation above.

Figure 28: Proof for the annotation of metabolite GE20372 A or B.

KF 77 AG6
Found: 366.1769
Calc: 366.1772
Δppm: 1

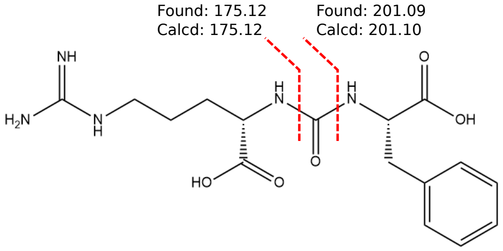

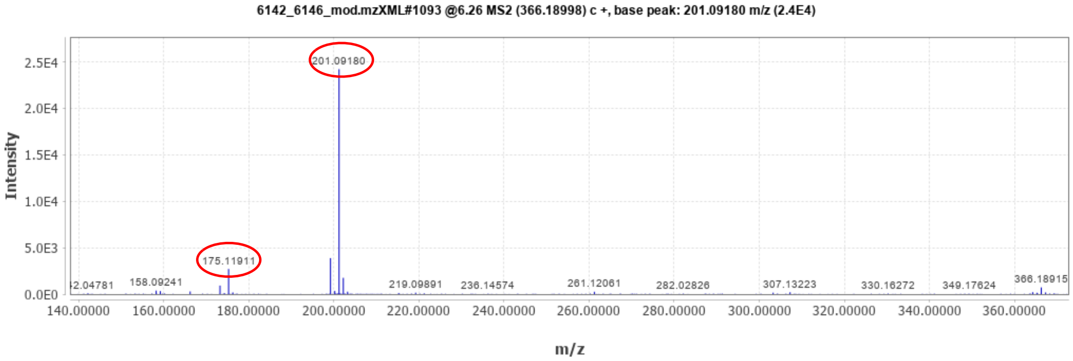

ESI-MS/MS spectrum for the [M+H]+ ion of **KG 77 AG6**, measured on a Bruker micrOTOF-Q III instrument. Peaks circled in red are consistent with the fragmentation above.

Figure 29: Proof for the annotation of metabolite KF-77-AG6.

Desferrioxamine B
Found: 561.36 m/z
Calc: 561.36 m/z
Δppm: 5.9

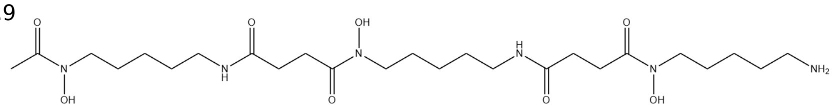

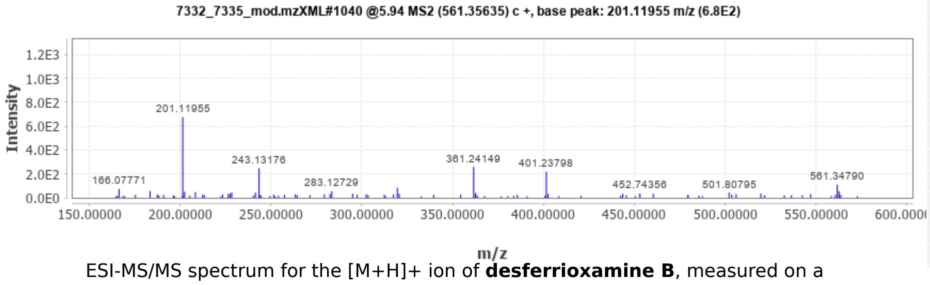

ESI-MS/MS spectrum for the [M+H]+ ion of **desferrioxamine B**, measured on a Bruker micrOTOF-Q III instrument, detected in the extracts 7332 and 7335.

B)

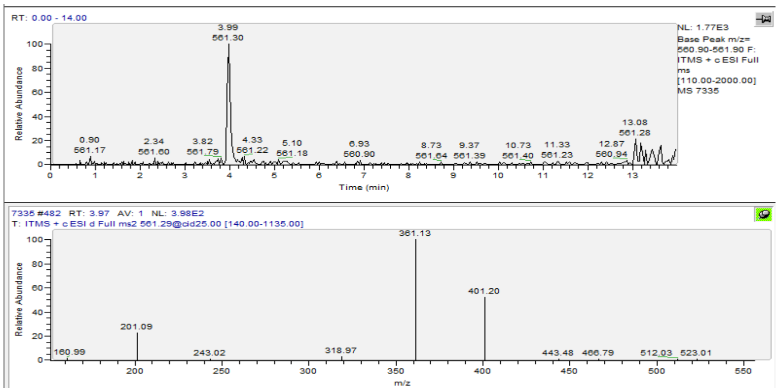

ESI-MS/MS spectrum for the [M+H]+ ion of **desferrioxamine B**, measured on a ThermoFisher LTQ linear ion trap instrument, detected in the extract **7335**. The main MS2 fragments correspond to the ones detected on the high resolution instrument (A). The retention time and the main MS2 fragments correspond to the ones detected for the Desferoxamine standard (C).

C)

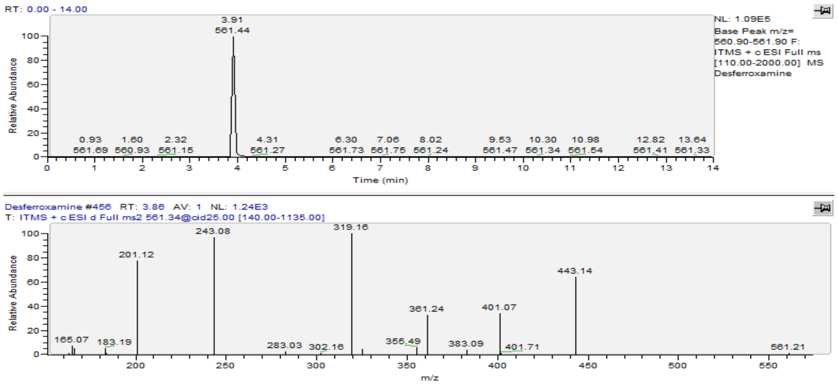

ESI-MS/MS spectrum for the [M+H]+ ion of desferrioxamine B, measured on a ThermoFisher LTQ linear ion trap instrument, from **Deferoxamine mesylate salt standard** of Sigma Aldrich (Prod. Nr. D9533, CAS 138-14-7).

Figure 30: Proof for the annotation of metabolite Desferrioxamine B.

97518/planosporicin
Found: 1096.8967
Calc: 1096.8902
Δppm: 6

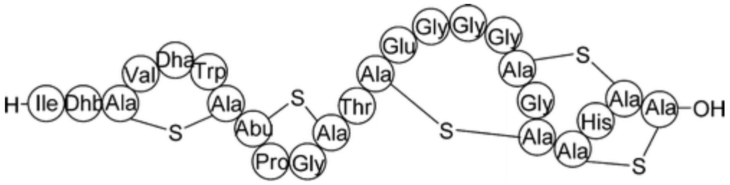

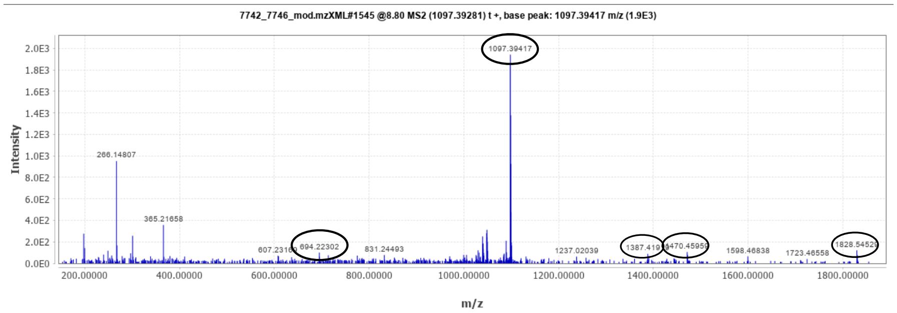

ESI-MS/MS spectrum for the [M+2H+1]2+ ion of **97518/planosporicin**, measured on a Bruker micrOTOF-Q III instrument. Peaks circled in black are consistent with the published spectrum (Maffioli et al. J Nat Prod 72.4 (2009) 605-607).

Figure 31: Proof for the annotation of metabolite lanthibiotic 97518/planosporicin.

Sphaericin
[M+H+1]+
Found: 2157.116
Calc: 2157.112
Δppm: 0.8

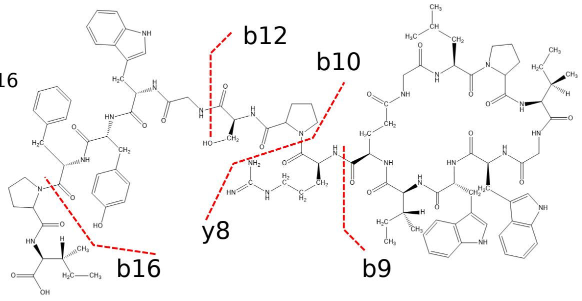

| Fragment | Found | Calcd |
|---|---|---|
| b16 | 1927.9651 | 1927.9649 |
| b12 | 1374.7277 | 1374.7322 |
| b10 | 1190.6306 | 1190.6474 |
| b9 | 1034.5548 | 1034.5463 |
| y8 | 966.4675 | 966.4725 |

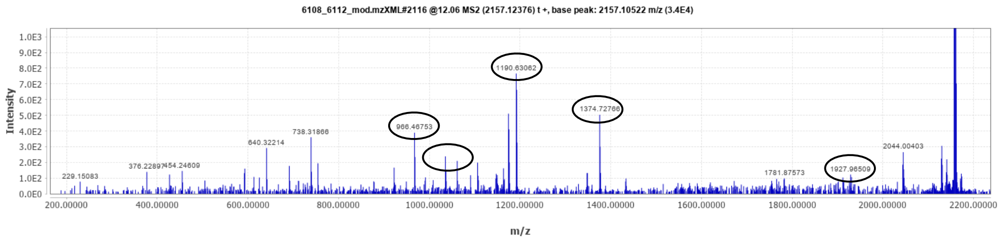

ESI-MS/MS spectrum for the [M+H+1]+ ion of **Sphaericin**, measured on a Bruker micrOTOF-Q III instrument. Peaks circled in black are consistent with the published spectrum (Kodani et al. Europ J Org Chem 2017.8 (2017): 1177-1183).

Figure 32: Proof for the annotation of metabolite sphaericin.

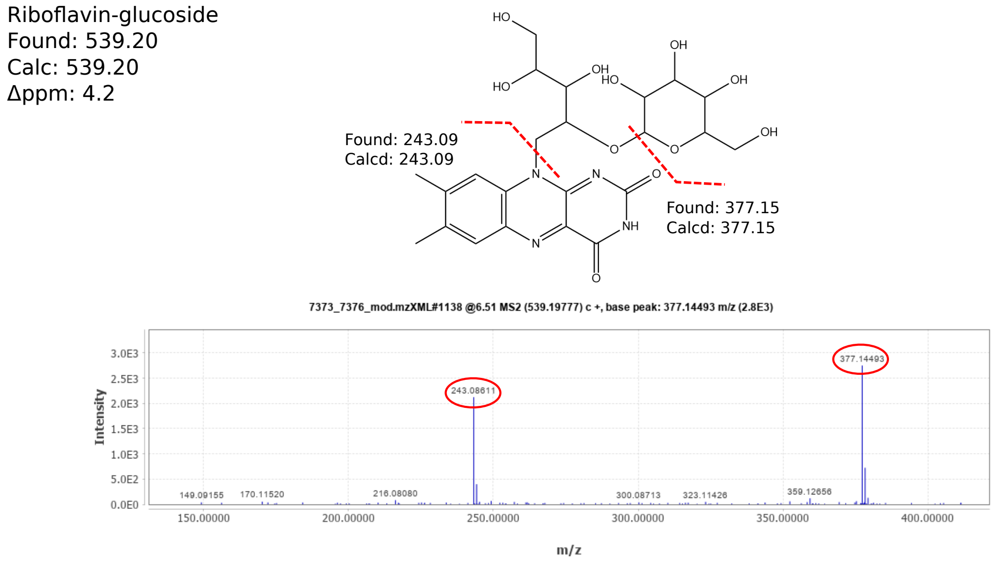

ESI-MS/MS spectrum for the [M+H]+ ion of r**iboflavin-glucoside**, measured on a Bruker micrOTOF-Q III instrument. Peaks circled in red are consistent with the fragmentation above.

Figure 33: Proof for the annotation of metabolite riboflavin-glucoside.

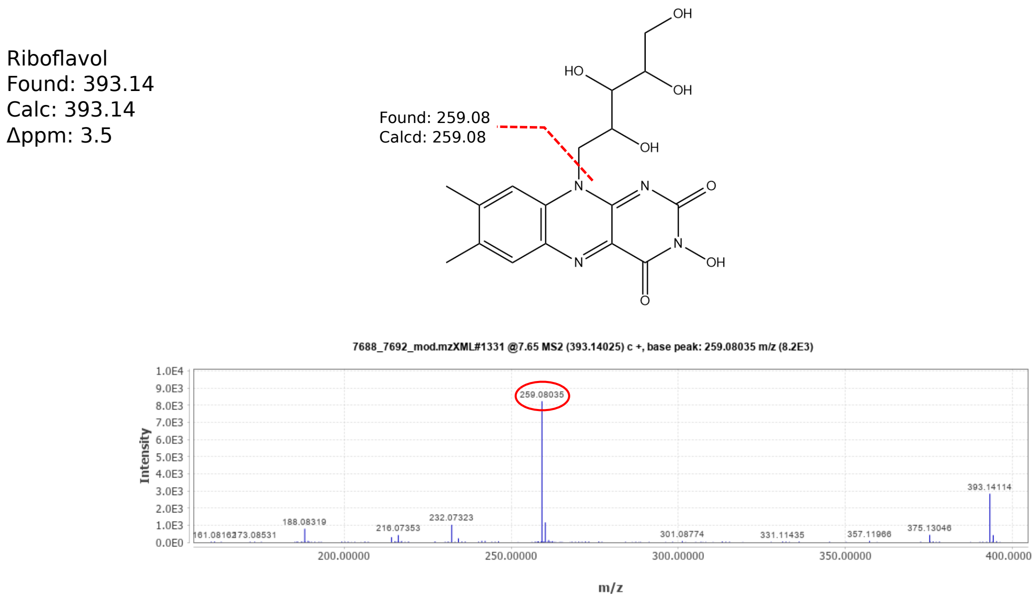

ESI-MS/MS spectrum for the [M+H]+ ion of r**iboflavol**, measured on a Bruker micrOTOF-Q III instrument. Peaks circled in red are consistent with the fragmentation above.

Figure 34: Proof for the annotation of metabolite riboflavol.

**Bruker DataAnalysis script**

```
option explicit

Dim Filename

Analysis.Compound.Clear

Analysis.ClearResults

Analysis.Save

Filename = left(Analysis.path,len(Analysis.path)-2) & ".mzXML"

Analysis.Export Filename, daMzXML, daLine

Analysis.FindAutoMSn

Filename = left(Analysis.path,len(Analysis.path)-2) & ".mgf"

Analysis.Compounds.Export Filename, daMGF

Form.Close
```

Automatic export of .mzXML and .mgf-files. For FindAutoMSn. the intensity threshold was set to 0 and max. nr. compounds to 100000.

Figure 35: Transcript of Bruker method-file for the bulk export of .mzXML and .mgf-files.

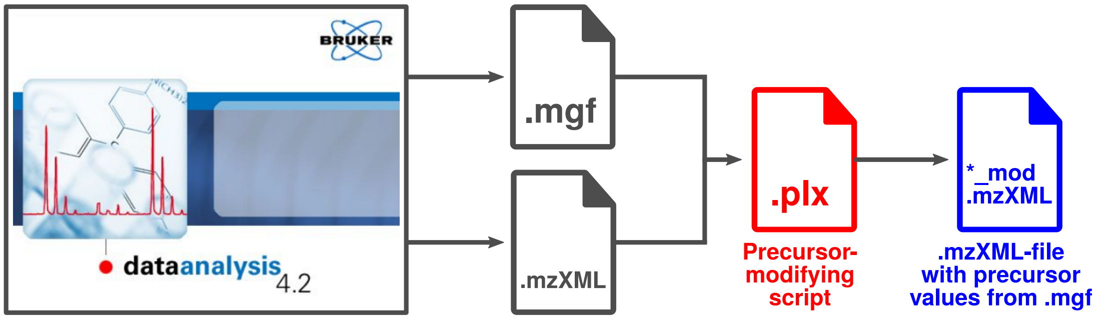

Figure 36: Visualisation of precursor correction workflow. Both .mzXML and .mgf-files are exported from the proprietary Bruker .raw-format using Bruker Data Analysis software. The Perl5 "precursor modification script" uses the correct precursor values from the .mgf-file to modify the un-calibrated precursor values from the .mzXML. A new file with the suffix "_mod.mzXML" is created and used in the consequent data processing.

**Perl (v5.22.1) precursor modification script**

```
#!/usr/bin/env perl
use strict;
use warnings;
use autodie;
use feature qw(say);
my %hash;
#opens .mgf-file (second argument in bash command)
open my $fhh, "<", "$ARGV[1]";
my $key = 0;
my $val = 0;
my $tag = 0;
#Processes .mgf-file: each entry (content between "BEGIN IONS" and "END IONS") contains a "SCANS" entry that gives information to the
#scans it comes from (e.g. "SCANS=MS: 317 MSMS: 320" means "MS2 scan #320, orginating from MS1 scan 317").The "PEPMASS"-field
#corresponds to the (correct) precursor-m/z. The first loop checks every line of the .mgf-file for the MS2-scan and assigns it to variable $key.
#This also sets the flag $tag which stores the precursor-value in the next line in a hash, using the MS2-id as key. The fact that the
#"PEPMASS" always follows "SCANS" gives specificity.
while (my $input = <$fhh>){
 chomp $input;
 if ( $input =~ m/\ASCANS=MS:\ [\d]+\ MSMS:\ ([\d]+)/ ) {
  $key = $1 if $input =~ m/\ASCANS=MS:\ [\d]+\ MSMS:\ ([\d]+)/;
  $tag = 1;
 }
 if ( $tag ) {
  $val = $1 if $input =~ m/\APEPMASS=([\d]+\.[\d]+)/;
  if ( $val ){
   $hash{$key} = $val;
   $key = 0;
   $val = 0;
   $tag = 0;
  }
 }
}
close $fhh;
#Opens .mzXML-file (accompanying the .mgf-file) and filehandle for output-file
open my $fh, "<", "$ARGV[0]";
open my $out, ">", "modified_$ARGV[0]";
my $flag = 0;
my $id = 0;
#Processes .mzXML-file and writes output to new file: each MS2-scan in a .mzXML-file starts with a header containing the scan-ID and a
#"msLevel="2"" fields, followed by a line containing information regarding the precursor-mz and intensity. The next loop checks each line for
#the "marker" msLevel="2". If encountered, the scan-id is compared to all keys in the previously created hash. If there is a match, a flag is set.
#In the next line, the erroneous precursor is replaced with the one extracted from the .mgf-file and the flags are reset. Again, specificity is
#given because the line containing the precursor is always precedet by the scan-id.
while (my $line = <$fh>) {
 chomp $line;
  if ( $line =~ m`msLevel=\"2\"` ) {
   foreach my $i (keys %hash) {
   next unless $line =~ m`\A(<scan\ num=\"$i\")`;
   $id = $i;
   $flag = 1;
   last;
   }
 }
 if ( $flag ) {
  if ( $line =~ m`\A<precursorMz\ precursorIntensity=` ) {
   $line =~ s`[\d]+\.[\d]+(</precursorMz>)`$hash{$id}$1`;
   $id = 0;
   $flag = 0;
  }
 }
print $out "$line\n";
}
close $fh;
close $out;
```

Figure 37: Script for replacing un-calibrated precursor values in .mzXML-files with the calibrated ones from the .mgf-file.

**Genes considered in autoMLST phylogenetic tree**

| Protein Family | Gene | Associated Function | Protein Family | Gene | Associated Function |
|---|---|---|---|---|---|
| TIGR00133 | gatB | Protein synthesis | TIGR01855 | IMP_synth_hisH | Amino acid biosynthesis |
| TIGR00132 | gatA | Protein synthesis | TIGR00174 | miaA | Protein synthesis |
| TIGR03953 | rplD_bact | Protein synthesis | TIGR00647 | DNA_bind_WhiA | Cellular processes |
| TIGR00138 | rsmG_gidB | Protein synthesis | TIGR01083 | nth | DNA metabolism |
| TIGR01520 | FruBisAldo_II_A | Energy metabolism | TIGR00577 | fpg | DNA metabolism |
| TIGR01529 | argR_whole | Regulatory functions | TIGR01169 | rplA_bact | Protein synthesis |
| TIGR02027 | rpoA | Transcription | TIGR00928 | purB | Purines, pyrimidines |
| TIGR01203 | HGPRTase | Purines, pyrimidines | TIGR01327 | PGDH | Amino acid biosynthesis |
| TIGR00337 | PyrG | Purines, pyrimidines | TIGR01994 | SUF_scaf_2 | Biosynthesis of cofactors etc. |
| TIGR00331 | hrcA | Regulatory functions | TIGR01164 | rplP_bact | Protein synthesis |
| TIGR01049 | rpsJ_bact | Protein synthesis | TIGR01009 | rpsC_bact | Protein synthesis |
| TIGR01044 | rplV_bact | Protein synthesis | TIGR00168 | infC | Protein synthesis |
| TIGR00962 | atpA | Energy metabolism | TIGR00042 | TIGR00042 | DNA metabolism |
| TIGR00963 | secA | Protein fate | TIGR00048 | rRNA_mod_RlmN | Protein synthesis |
| TIGR01966 | RNasePH | Transcription | TIGR00281 | TIGR00281 | DNA metabolism |
| TIGR00459 | aspS_bact | Protein synthesis | TIGR00184 | purA | Purines, pyrimidines |
| TIGR00755 | ksgA | Protein synthesis | TIGR00166 | S6 | Protein synthesis |
| TIGR00521 | coaBC_dfp | Biosynthesis of cofactors etc. | TIGR00713 | hemL | Biosynthesis of cofactors etc. |
| TIGR03594 | GTPase_EngA | Siomycin-related motif | TIGR02274 | dCTP_deam | Purines, pyrimidines |
| TIGR00615 | recR | Protein synthesis | TIGR00362 | DnaA | DNA metabolism |
| TIGR00244 | TIGR00244 | Regulatory functions | TIGR03654 | L6_bact | Protein synthesis |
| TIGR01039 | atpD | Energy metabolism | TIGR01171 | rplB_bact | Protein synthesis |
| TIGR01032 | rplT_bact | Protein synthesis | TIGR00959 | ffh | Protein fate |
| TIGR00019 | prfA | Protein synthesis | TIGR01455 | glmM | Central intermediary metabolism |
| TIGR01978 | sufC | Biosynthesis of cofactors etc. | TIGR01071 | rplO_bact | Protein synthesis |
| TIGR02673 | FtsE | Cellular processes | TIGR00952 | S15_bact | Protein synthesis |
| TIGR00150 | T6A_YjeE | Protein synthesis | TIGR00119 | acolac_sm | Amino acid biosynthesis |
| TIGR00420 | trmU | Protein synthesis | TIGR00461 | gcvP | Energy metabolism |
| TIGR00090 | rsfS_iojap_ybeB | Protein synthesis | TIGR00510 | lipA | Biosynthesis of cofactors etc. |
| TIGR00096 | TIGR00096 | Protein synthesis | TIGR00355 | purH | Purines, pyrimidines |
| TIGR01302 | IMP_dehydrog | Purines, pyrimidines | TIGR01736 | FGAM_synth_II | Purines, pyrimidines |
| TIGR03800 | PLP_synth_Pdx2 | Biosynthesis of cofactors etc. | TIGR01737 | FGAM_synth_I | Purines, pyrimidines |
| TIGR01021 | rpsE_bact | Protein synthesis | TIGR01066 | rplM_bact | Protein synthesis |
| TIGR01024 | rplS_bact | Protein synthesis | TIGR01063 | gyrA | DNA metabolism |
| TIGR00065 | ftsZ | Cellular processes | TIGR02692 | tRNA_CCA_actino | Protein synthesis |
| TIGR00064 | ftsY | Protein fate | TIGR02729 | Obg_CgtA | Protein synthesis |
| TIGR00060 | L18_bact | Protein synthesis | TIGR00020 | prfB | Protein synthesis |
| TIGR01394 | TypA_BipA | Regulatory functions | TIGR00580 | mfd | DNA metabolism |
| TIGR00436 | era | Protein synthesis | TIGR00228 | ruvC | DNA metabolism |
| TIGR00635 | ruvB | DNA metabolism | TIGR00670 | asp_carb_tr | Purines, pyrimidines |
| TIGR00302 | TIGR00302 | Purines, pyrimidines | TIGR01059 | gyrB | DNA metabolism |
| TIGR00431 | TruB | Protein synthesis | TIGR00343 | TIGR00343 | Biosynthesis of cofactors etc. |
| TIGR01980 | sufB | Biosynthesis of cofactors etc. | TIGR03631 | uS13_bact | Protein synthesis |
| TIGR01011 | rpsB_bact | Protein synthesis | TIGR01051 | topA_bact | DNA metabolism |
| TIGR00468 | pheS | Protein synthesis | | | |

Figure 38: Genes considered for the multilocus sequence analysis (autoMLST).
